# Endogenous LRRK2 and PINK1 function in a convergent neuroprotective ciliogenesis pathway in the brain

**DOI:** 10.1101/2024.06.11.598416

**Authors:** Enrico Bagnoli, Yu-En Lin, Sophie Burel, Ebsy Jaimon, Odetta Antico, Christos Themistokleous, Jonas M. Nikoloff, Ilaria Morella, Jens O. Watzlawik, Fabienne C. Fiesel, Wolfdieter Springer, Francesca Tonelli, Simon P. Brooks, Stephen B. Dunnett, Riccardo Brambilla, Dario R. Alessi, Suzanne R. Pfeffer, Miratul M. K. Muqit

## Abstract

Mutations in LRRK2 and PINK1 are associated with familial Parkinson’s disease (PD). LRRK2 phosphorylates Rab GTPases within the Switch II domain whilst PINK1 directly phosphorylates Parkin and ubiquitin and indirectly induces phosphorylation of a subset of Rab GTPases. Herein we have crossed LRRK2 [R1441C] mutant knock-in mice with PINK1 knock-out (KO) mice and report that loss of PINK1 does not impact endogenous LRRK2-mediated Rab phosphorylation nor do we see significant effect of mutant LRRK2 on PINK1-mediated Rab and ubiquitin phosphorylation. In addition, we observe that a pool of the Rab-specific, PPM1H phosphatase, is transcriptionally up-regulated and recruited to damaged mitochondria, independent of PINK1 or LRRK2 activity. Parallel signalling of LRRK2 and PINK1 pathways is supported by assessment of motor behavioural studies that show no evidence of genetic interaction in crossed mouse lines. Previously we showed loss of cilia in LRRK2 R1441C mice and herein we show that PINK1 KO mice exhibit a ciliogenesis defect in striatal cholinergic interneurons and astrocytes that interferes with Hedgehog induction of glial derived-neurotrophic factor (GDNF) transcription. This is not exacerbated in double mutant LRRK2 and PINK1 mice. Overall, our analysis indicates that LRRK2 activation and/or loss of PINK1 function along parallel pathways to impair ciliogenesis, suggesting a convergent mechanism towards PD. Our data suggests that reversal of defects downstream of ciliogenesis offers a common therapeutic strategy for LRRK2 or PINK1 PD patients whereas LRRK2 inhibitors that are currently in clinical trials are unlikely to benefit PINK1 PD patients.

## INTRODUCTION

Gain-of-function mutations in Leucine-rich repeat kinase 2 (LRRK2) are associated with autosomal dominant Parkinson’s disease (PD) and are frequently found in sporadic PD patients [1–3]. LRRK2 encodes a multidomain protein containing both a guanosine triphosphatase (GTPase) [Ras of complex (Roc)-C-terminal of Roc (COR)] domain and a protein kinase domain, together with N-terminal protein-interaction domains, recently shown to bind Rab GTPases at distinct sites that regulate the activation state of LRRK2 at membranes [4–7]. Pathogenic missense mutations span all domains but are predominantly located within the kinase (e.g. [Gly2019Ser (G2019S)] or Roc-COR domains (e.g. [Arg1441Cys/Gly/His (R1441C/G/H)] and lead to elevation of LRRK2 kinase activity [8]. LRRK2 phosphorylates a subset of Rab GTPases, including Rab1, Rab3. Rab8, Rab10, Rab12, Rab29, Rab35, and Rab43, at a highly conserved Ser/Thr residue positioned within the Switch II-effector binding domain (Rab8 Thr72; Rab10 Thr73; and Rab12 Ser106) [9, 10]. LRRK2-mediated phosphorylation of Rab proteins stimulates binding to RILPL1/2 and JIP3/4 proteins that regulate downstream processes including ciliogenesis [9, 11, 12] and lysosomal stress responses [13–15]. Conversely, the PPM1H phosphatase dephosphorylates the Switch II domain-phosphorylated residues of LRRK2-regulated Rabs and this has been shown to counteract LRRK2’s effects on phenotypes such as primary ciliogenesis [16, 17].

Loss-of-function mutations in PTEN-induced kinase 1 (PINK1) cause autosomal recessive PD [18]. PINK1 contains an N-terminal mitochondrial targeting domain and a protein kinase domain, flanked by N-terminal and C-terminal regions that facilitate recruitment of PINK1 to the Translocase of outer membrane (TOM) complex that is required for its activation at sites of mitochondrial damage [19–23]. Most of pathogenic mutations are located within the kinase domain and disrupt its catalytic activity [24–26]. Active PINK1 directly phosphorylates Parkin at a conserved Ser65 residue that lies within its ubiquitin-like domain (Ubl) and an equivalent Ser65 residue on ubiquitin, resulting in activation of Parkin via a feed-forward mechanism, triggering ubiquitin-dependent elimination of damaged mitochondria by autophagy (mitophagy) [27–34]. Active PINK1 also indirectly induces the phosphorylation of a subset of Rab GTPases including Rab1, Rab8 and Rab13 at a highly conserved Ser residue located within the RabSF3 effector-binding motif (Rab8 Ser111), distinct from the site modified by LRRK2 [35–37]. PINK1-mediated Rab phosphorylation inhibits binding of effector proteins including guanine exchange factor (GEF) and GTPase activating proteins (GAP) [35–37]. We previously reported that PINK1 phosphorylation of Rab8 Ser111 impairs the ability of LRRK2 to phosphorylate Thr72 *in vitro,* however, whether this occurs in cells under endogenous protein expression conditions has hitherto not been assessed [36].

Previous studies have reported cross-talk between LRRK2 and PINK1 mitophagy signalling. An early report suggested that LRRK2 protein expression is increased in human fibroblasts and induced pluripotent stem cell (iPSC)-derived dopaminergic neurons derived from compound heterozygous deletion mutant or homozygous G309D PINK1 mutant patients [38]. A further study showed that PINK1-dependent mitochondrial depolarisation-induced mitophagy is impaired in human fibroblasts derived from PD patients harbouring the LRRK2 G2019S or R1441C mutations and this could be rescued by LRRK2 genetic knockdown or inhibitor treatment [39]. The authors suggested this was mediated by LRRK2-mediated Rab10 phosphorylation that impaired its interaction with the autophagy receptor, Optineurin at mitochondria [39]. It has also been reported that PINK1-dependent mitophagy was impaired by hyperactive LRRK2 mutations in cell lines and human patient-derived LRRK2 [G2019S] fibroblasts [40]. A more recent study reported that decreased mitochondrial depolarisation-induced mitophagy in primary cortical neurons derived from LRRK2 R1441C transgenic rats is associated with a significant decrease in phosphorylated ubiquitin compared to non-transgenic neurons [41]. That study also showed significant decrease in mitochondrial depolarisation-induced accumulation of phosphorylated ubiquitin and concomitant mitophagy in human iPSC-derived dopaminergic neurons derived from human fibroblasts expressing the LRRK2 R1441C mutation, although this was not rescued by the LRRK2 inhibitor, MLi-2 [41]. Furthermore, LRRK2 has been implicated in regulation of basal mitophagy and analysis of mitophagy in G2019S mutant mice *in vivo* revealed higher levels of basal mitophagy that was independent of PINK1 and could also be rescued by LRRK2 inhibitors [42, 43].

In this study we have assessed whether there is any physiological regulation of endogenous LRRK2 by endogenous PINK1 and vice versa in mouse tissues and mouse embryonic fibroblasts (MEFs). We have generated double mutant PINK1/LRRK2 mice models to assess the role of PINK1 on basal wild-type LRRK2 and pathogenic mutant [R1441C] activity, and conversely, the impact of LRRK2 hyperactivation on PINK1 basal activity in mitochondrial depolarisation-dependent activity in MEFs. Our data indicate that knock-out of PINK1 does not impact the ability of LRRK2 to phosphorylate its Rab substrates; similarly, we do not observe any significant effect of LRRK2 activity on endogenous PINK1-dependent substrate phosphorylation. However, we report for the first time a downstream role for PINK1 in the regulation of cilia and GDNF production in the striatum.

## RESULTS

### PINK1 knock-out does not impact basal LRRK2-mediated phosphorylation of Rab12 and Rab10 in mouse brain and peripheral tissues

To investigate the role of endogenous PINK1 in regulating LRRK2 signalling, we crossed LRRK2 R1441C knock-in mice [44–47] with PINK1 knock-out (KO) mice [48–50] that we have characterised in previous studies (Figure 1A). Double mutant LRRK2 [R1441C] / PINK1 KO mice were viable and displayed no overt phenotypes. As a readout for LRRK2 pathway activity, we measured phosphorylation of Rab12 at Ser105 (equivalent phosphorylation site to human pSer106) and Rab10 at Thr73, using well-characterised phospho-specific antibodies [14, 44]. Immunoblot analysis of sub-dissected striatal (Figure 1 B-F) and cortical (Figure 1 G-K) brain regions from mice treated with or without the LRRK2 inhibitor, MLi-2, confirmed previous data in whole brain that the LRRK2 [R1441C] mutation enhances LRRK2-mediated Rab12 and Rab10 phosphorylation (Figure 1 B, D and G, I) and decreases LRRK2 Ser935 phosphorylation (Figure 1 B, E and G, J) [14, 47]. Furthermore, LRRK2-phosphorylated Rab12 or Rab10, quantified in relation to total Rab protein, was not affected by PINK1 knock-out (Figure 1 B, D and G, I). As expected, MLi-2 treatment markedly reduced Rab12 and Rab10 phosphorylation, concomitant with a decrease in LRRK2 Ser935 phosphorylation (Figure 1 B, E and G, J). We also did not observe any significant changes in the total levels of LRRK2 or PPM1H in PINK1 KO mice lines (Figure 1 B, F and G, K). Broadly similar results were observed for other brain regions analysed including olfactory bulb (Supplementary Figure 1 A, C-E); hippocampus (Supplementary Figure 1 B, F-H), midbrain (Supplementary Figure 2 A, C-E), thalamus (Supplementary Figure 2 B, F-H), cerebellum (Supplementary Figure 3 A, C-E), brainstem (Supplementary Figure 3 B, F-H), and spinal cord (Supplementary Figure 4 A-D).

**Figure 1.**
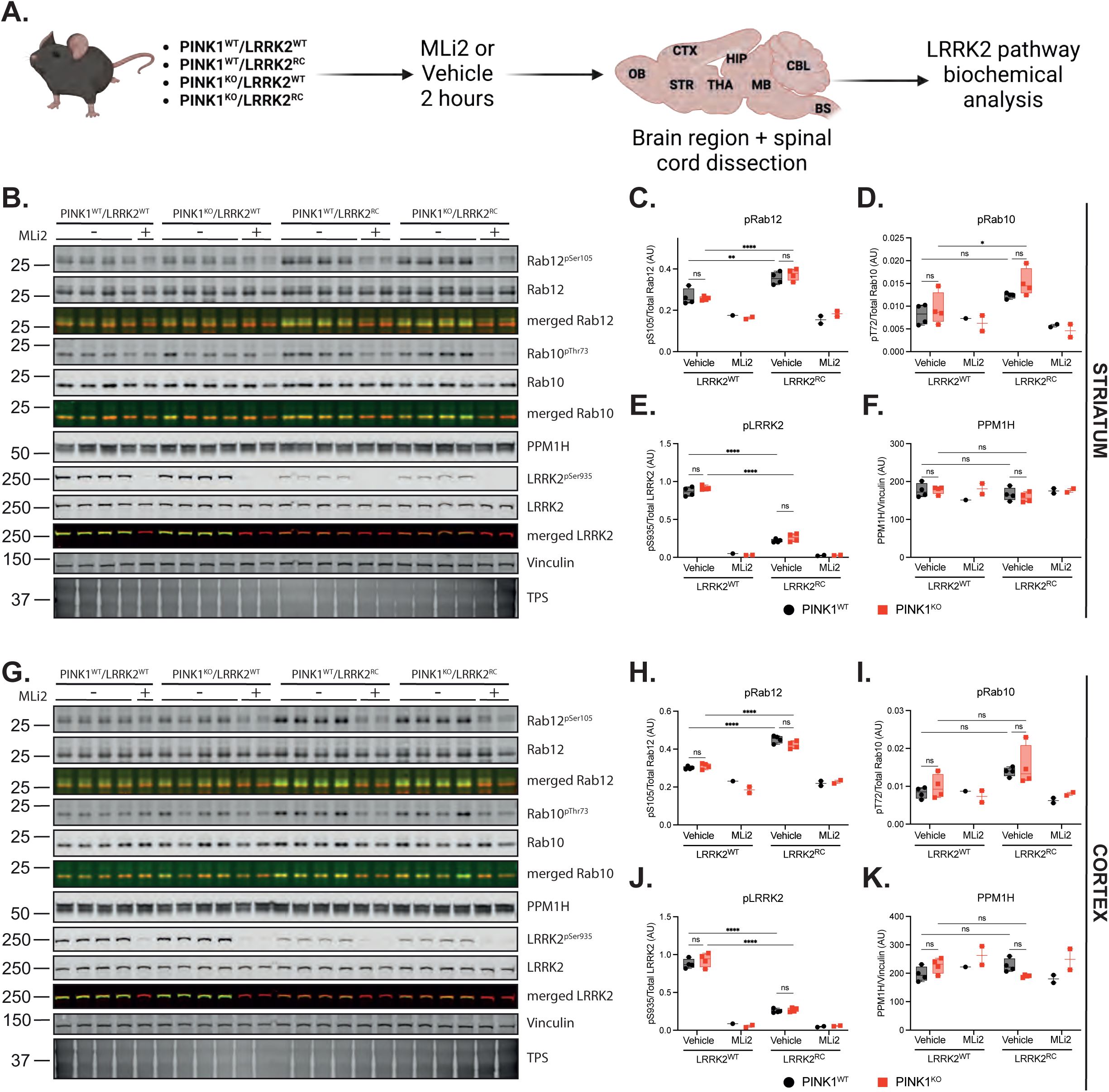
LRRK2 signalling pathway in the midbrain and cortex is not affected by loss of PINK1 *in vivo*. ***A.*** Schematic of methodology followed ***B.*** Immunoblot of LRRK2 pathway component in mouse striatum and relative quantification of ***C.*** pSer105/total Rab12, ***D.*** pThr73/total Rab10, ***E.*** pSer935/total LRRK2, and ***F.*** PPM1H/Vinculin. Similarly in ***G., H., I., J., K.*** analysis from mouse cortex. Each lane was loaded with 40 μg of protein lysate from one mouse. In graphs, black circle represents PINK1^WT^ while red square PINK1^KO^ animals. Box and whiskers plot, from min to max with the median line. Ordinary 2-way ANOVA with Sidak’s multiple comparison test. * p<0.05, ** p<0.01, *** p<0.001, **** p<0.0001.

In parallel experiments, we analysed LRRK2 signalling in peripheral mouse tissues including lung and spleen (Supplementary Figure 5). LRRK2-phosphorylated Rab10 or Rab12, quantified in relation to total Rab protein, were not affected by PINK1 knock-out in lung (Supplementary Figure 5 A-C). Similarly, levels of LRRK2 Ser935 phosphorylation and total PPM1H expression were unchanged between WT or PINK1 KO mice (Supplementary Figure 5A, D, E). Furthermore, we observed no impact of PINK1 knock-out on LRRK2-phosphorylated Rab10 levels in the spleen or on LRRK2 Ser935 phosphorylation (Supplementary Figure 5 F-I). Total Rab12 levels in the spleen were low, which prevented assessment of LRRK2-phosphorylated Rab12.

### Behavioural motor analysis of aged mice does not indicate interplay between LRRK2 and PINK1 pathways *in vivo*

Previous analyses of LRRK2 [R1441C] knock-in mice and PINK1 KO mice has not detected an overt Parkinsonian motor phenotype [42–48] although we have recently found decreased striatal dopamine projections and GDNF Receptor alpha staining in LRRK2 mutant mice models [51]. We did not observe any gross difference in weight across all four mouse genotypes of 18 wild-type, LRRK2 [R1441C], PINK1 knock-out (KO), or the double mutant LRRK2 [R1441C] / PINK1 KO mice (Figure 2 B and Supplementary Figure 6 G-H). We next quantitatively characterised motor function using three of the most widely used behavioural tests, namely rotarod, balance beam, and gait analysis [52, 53] (Figure 2 A). For rotarod testing we observed slight reduction in the latency to fall for LRRK2 [R1441C] mice compared with wild-type control mice but this was not significantly altered in double mutant LRRK2 [R1441C] / PINK1 KO mice (Figure 2 G). For the balance beam, we observed a slight increase in the latency to turn by LRRK2 [R1441C] mice compared with wild-type control mice (Figure 2 F), associated with a slight increase in the number of forelimb and hindlimb slips (Supplementary Figure 6 D-F), however, similar to rotarod testing, this was not significantly altered in double mutant LRRK2 [R1441C] / PINK1 KO mice (Figure 2 F and Supplementary Figure 6 D-F). Interestingly, gait analysis revealed a subtle decrease in stride length of the PINK1 KO mice, probably due to the smaller dimension of the mice; however, this was not significantly different in the double mutant LRRK2 [R1441C] / PINK1 KO mice (Figure 2 E). Overall, the motor impairments observed were selective and subtle, and consistent with this we did not observe any impairment for other measures of gait analysis including width of forelimb or hindlimb base (Supplementary Figure 6 A-C) nor in grip strength (Figure 2 D) or proprioception (Figure 2 C). Immunohistochemical analysis of brain sections did not reveal any difference in DARPP-32 staining of striatal medium spiny neurons or total striatal volume between mice of different genotypes (Supplementary Figure 7 A-C). However, analysis of microglia revealed an increase in their number in LRRK2 [R1441C] and PINK1 KO animals, but this was not further exacerbated in the double mutant (Figure 2 H-I).

**Figure 2.**
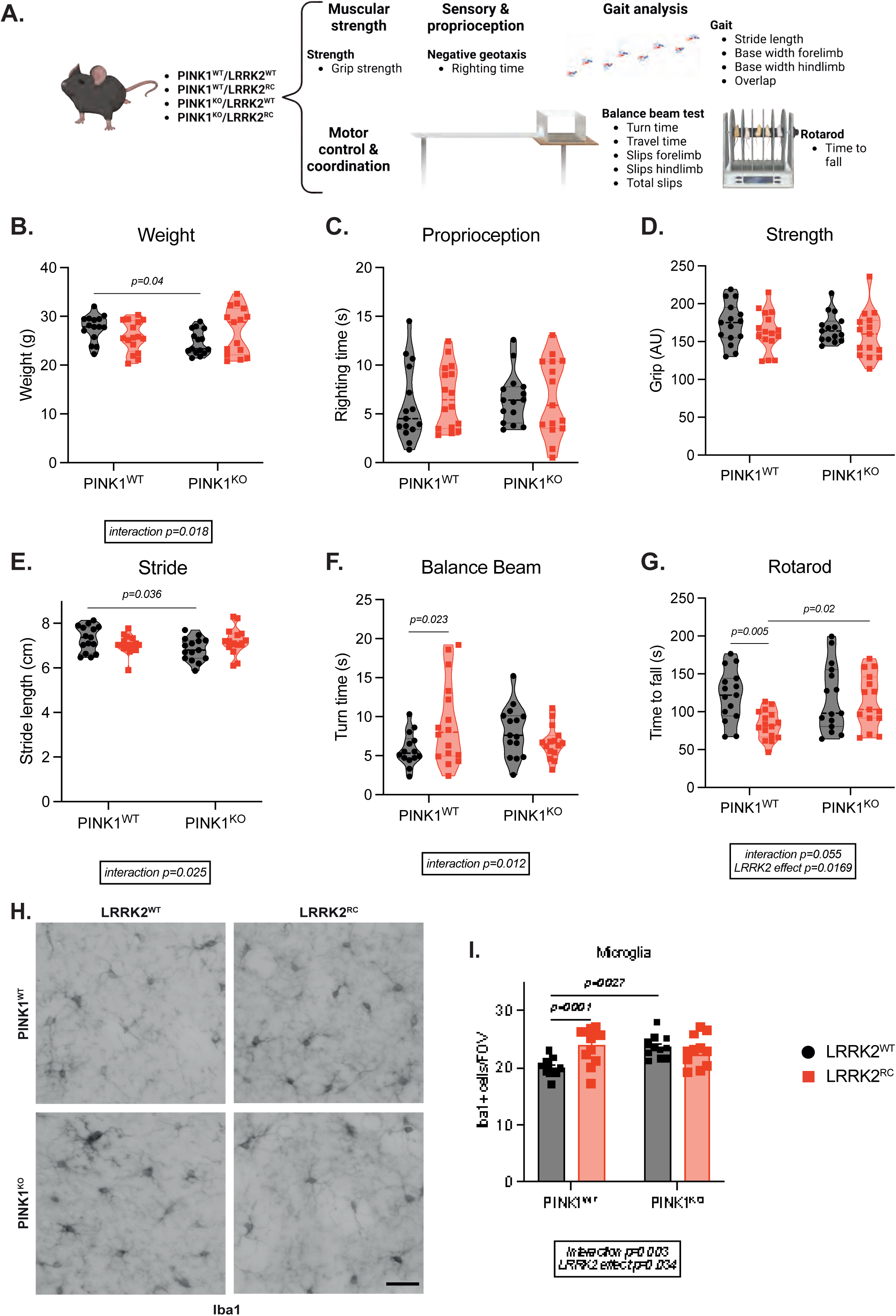
Behavioural testing in 10.5 months old double mutant PINK1 knockout / LRRK2 R1441C mutant mice does not suggest genetic interaction *in vivo*. ***A.*** A battery of different tests was performed to assess motor function in PINK1 knockout (KO) and LRRK2 [R1441C] knockin mice (LRRK2^RC^). ***B.*** Weight at 10.5 months. ***C.*** Righting time from negative geotaxis test and ***D.*** grip strength. ***E.*** measure of stride length during gait analysis and time to turn ***F***. and time to fall ***G.*** from balance beam and rotarod tests. ***H.*** Representative images of the microglial marker Iba1 and ***I.*** quantification of the Iba1 positive cells. In violin plots, black circle represents LRRK2^WT^ while red squares LRRK2^RC^ mice. Ordinary 2-way ANOVA with Sidak’s multiple comparison test. * p<0.05, ** p<0.01, *** p<0.001, **** p<0.0001. N=15/16 mice per group. Scale bar 50 µm.

### PINK1 signalling pathway is not significantly impacted by mutant LRRK2 [R1441C]

We next investigated whether mutant LRRK2 impacts endogenous PINK1 signalling *in vivo*. It has recently been demonstrated that PINK1-dependent, phosphorylated ubiquitin is detectable in mouse tissues, including brain, under basal conditions, using an ELISA-based assay [54]. We therefore prepared sub-dissected brain regions (cortex, midbrain and cerebellum) and spinal cord from wild-type, LRRK2 [R1441C], PINK1 KO, or the double mutant LRRK2 [R1441C] / PINK1 KO mice and these were analysed by an independent laboratory in a blinded manner (Supplementary Figure 8). We did not observe any phosphorylated ubiquitin in samples obtained from the PINK1 KO or double mutant mice (Supplementary Figure 8 A-D). Overall we did not observe any significant difference in phosphorylated ubiquitin in select brain regions or spinal cord from LRRK2 [R1441C] mice compared to wild-type littermate control mice (Supplementary Figure 8 A-D), although interestingly there was a non-significant increase in phosphorylated ubiquitin in the midbrain of LRRK2 [R1441C] mice (Supplementary Figure 8 B).

Much of our understanding of PINK1 activation has been obtained under paradigms of mitochondrial damage which cannot be easily recapitulated *in vivo* [27, 28, 34]. Therefore, we next investigated whether LRRK2 activity influenced PINK1 mediated ubiquitin phosphorylation using wild-type and homozygous mutant LRRK2 R1441C MEFs treated with or without oligomycin and antimycin A (O/A) for 24 h to induce mitochondrial depolarisation in the presence or absence of PINK1 siRNA-mediated knockdown (Figure 3). Cells were lysed and whole cell extracts analysed by immunoblotting with anti-LRRK2 antibodies that confirmed uniform expression across all conditions (Figure 3 A). Following O/A treatment, we observed robust induction of phosphorylated ubiquitin and the PINK1-dependence was confirmed by loss of signal following siRNA-mediated PINK1 knock-down (Figure 3 A, B). Under these conditions we observed a non-significant mild increase in phosphorylated ubiquitin between homozygous LRRK2 R1441C mutant MEFs and wild-type controls (Figure 3 A, B), in line with the ELISA basal midbrain data. We further observed elevated basal phosphorylation of Rab10 at Thr73 (Figure 3 A, D) and Rab12 at Ser106 (Figure 3 A, E) in LRRK2 R1441C MEFs compared to wild-type controls and the LRRK2-dependence was confirmed by complete loss of phosphorylation following treatment with the LRRK2 inhibitor MLi-2 (Figure 3 A, D, E). We did not observe significant change in Rab 10 or 12 phosphorylation in the LRRK2 R1441C MEFs following O/A treatment in the presence or absence of PINK1 siRNA-mediated knockdown (Figure 3 A, D, E) indicating that LRRK2-mediated phosphorylation of Rabs is unaffected under conditions of PINK1 activation and consistent with the *in vivo* tissue analysis (Figure 1 and Supplementary Figures 1-5). Strikingly, we observed that PPM1H was up-regulated following O/A treatment in wild-type MEFs and this was similarly increased in LRRK2 R1441C mutant MEFs. Furthermore, this was not altered by siRNA-mediated knockdown of PINK1 (Figure 3 A and F).

**Figure 3.**
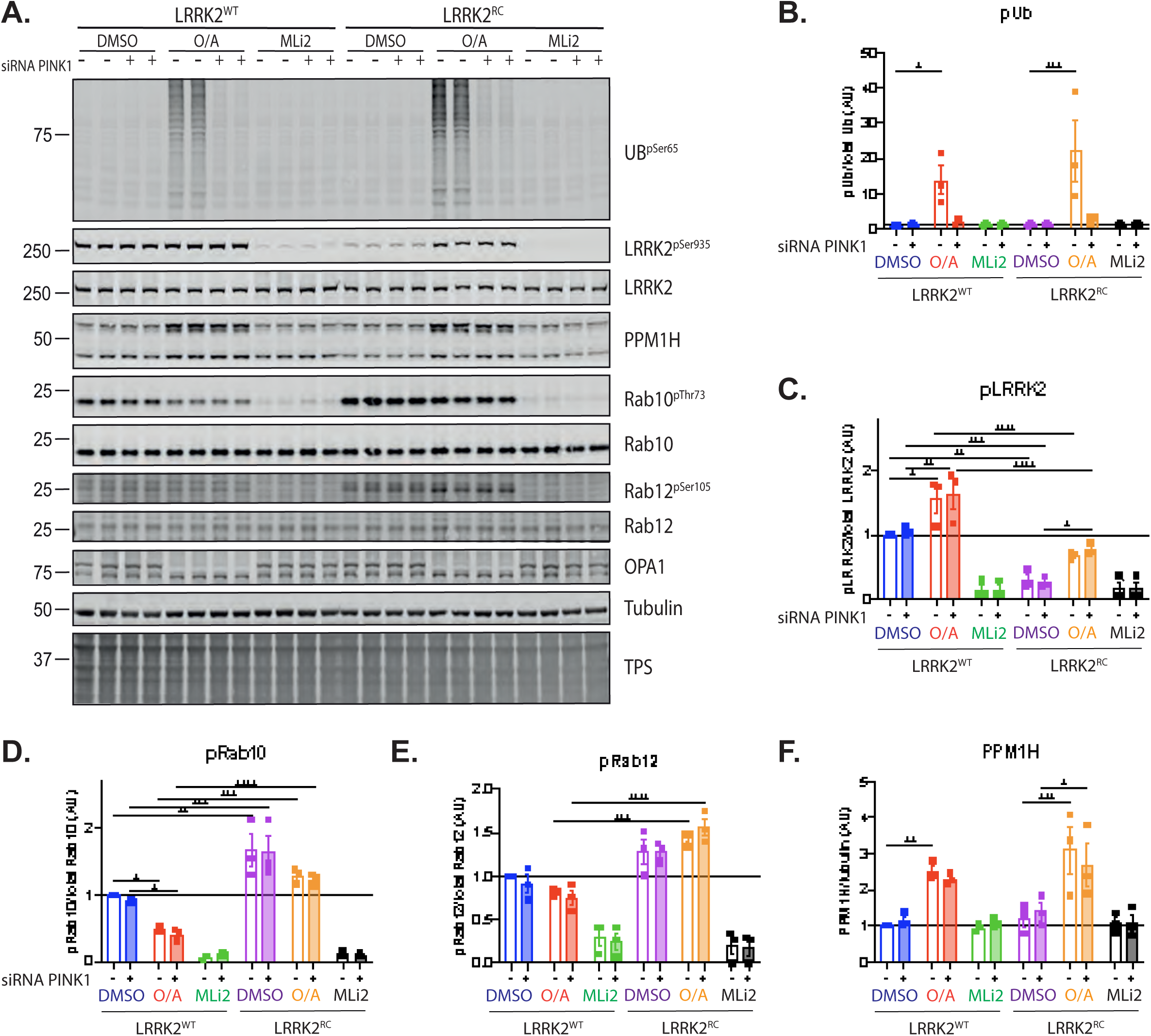
LRRK2 signalling pathway is not affected by mitochondrial damage-induced activation of PINK1. PINK1 was activated with oligomycin/antimycin for 24 hours, while LRRK2 was inhibited by 1 hour and 30 min MLi2 treatment ***A.*** Representative immunoblot of PINK1 activation effect on LRRK2 pathway components in LRRK2^WT^ and LRRK2^RC^ MEFs. Quantification from three experimental replicates for pUb, pLRRK2, pRab10, pRab12 and PPM1H is presented in ***B., C., D., E.,*** and ***F.*** respectively. Each lane was loaded with 30 μg of protein lysates from one well of a 6 well-plate. In graphs, each dot represents one experimental replicate with two biological replicates. - denotes scramble siRNA while + PINK1 siRNA. Bar graphs show mean ± SEM from three independent experiments. Ordinary 2-way ANOVA with Sidak’s multiple comparison test. * p<0.05, ** p<0.01, *** p<0.001, **** p<0.0001.

We next investigated whether LRRK2 activity influences endogenous PINK1-mediated Rab8A Ser111 phosphorylation. We therefore treated independent MEF clones derived from wild-type, LRRK2 [R1441C], PINK1 KO, or the double mutant LRRK2 [R1441C] / PINK1 KO mice, with or without O/A for 24 h to induce mitochondrial depolarisation and with MLi-2 to inhibit LRRK2 kinase activity (100nM for 1.5h, Supplementary Figure 9). Cells were lysed and whole cell extracts analysed by immunoblotting with anti-LRRK2 and GAPDH antibodies that confirmed uniform expression across all cell types and conditions (Supplementary Figure 9 A). We performed immunoprecipitation-based immunoblotting of total endogenous Rab8A followed by immunoblotting to detect Rab8A phosphorylated at Ser111 or at Thr72; and following O/A treatment, we observed robust induction of Ser111 Rab8A phosphorylation in wild-type MEF clones and observed no difference in both homozygous LRRK2 [R1441C] MEF clones tested (Supplementary Figure 9 A, B). In keeping with the PINK1-dependence of Ser111 Rab8A phosphorylation, we did not observe any signal in PINK1 KO or double mutant MEF clones (Supplementary Figure 9 A, B). Overall, these results suggest that endogenous PINK1-dependent activation and phosphorylation of Rab8A is not impacted by hyperactivation of endogenous LRRK2 catalytic activity. Consistent with previous analysis we observed basal LRRK2-mediated Thr72 Rab8A phosphorylation in wild-type MEFs and this was increased in homozygous LRRK2 [R1441C] MEF clones but unchanged in either O/A-treated cells or the double mutant LRRK2 [R1441C] / PINK1 KO MEFs (Supplementary Figure 9 A, C). These data further indicate that LRRK2 activity is not affected by basal or mitochondrial-induced activation of PINK1.

### PPM1H is up-regulated and recruited to damaged mitochondria

To investigate the up-regulation of PPM1H by O/A treatment, we employed previously generated homozygous PPM1H knock-out (KO) MEFs and corresponding wild-type control MEF clones [16, 55] and these were treated with O/A for 24 h in the presence or absence of MLi-2. Cells were lysed and whole cell extracts analysed by immunoblotting with anti-LRRK2 antibodies that confirmed uniform expression across all conditions (Supplementary Figure 10 A). Immunoblotting confirmed PPM1H up-regulation with O/A in wild-type MEFs and the signal was abolished in PPM1H KO MEFs (Supplementary Figure 10 A). Interestingly, the increase in PPM1H did not lead to any change/reduction in phosphorylated Rab10 (Supplementary Figure 10 A). In the presence of O/A, treatment with MLi-2 abolished phosphorylated Rab10 but did not impact the elevated PPM1H level (Supplementary Figure 10 A), pointing to a mechanism independent of LRRK2 kinase activity. Consistent with this, we did not observe any alteration of mitochondrial-stress induced PPM1H up-regulation in LRRK2 KO MEFs compared to respective wild-type controls (Supplementary Figure 10 B). In line with previous siRNA mediated PINK1 knockdown in MEFs (Figure 3 A), we also did not detect any changes in PPM1H up-regulation in PINK1 KO MEFs compared to wild-type controls (Supplementary Figure 10 C).

We next performed time course studies of PPM1H protein and mRNA expression following O/A treatment in wild-type MEFs by western blot and quantitative RT-PCR respectively (Figure 4). This revealed marked time-dependent increase of PPM1H protein evident at 16 h of O/A treatment (Figure 4 A, B) associated with minor changes in pLRRK2 and pRab10 (Supplementary Figure 11 A, B). Consistent with this we observed a significant increase in PPM1H mRNA from 4 h becoming maximal at 16 h using two independent primer pairs (Figure 4 C and Supplementary Figure 11 C) and under these conditions we also observed O/A-induced increase in ATF4 mRNA that was maximal at 4 h but sustained to 24 h (Supplementary Figure 11 D) as previously reported [56, 57]. To confirm O/A-induced transcriptional up-regulation of PPM1H, we used the transcriptional inhibitor 5,6-dichlorobenzimidazole 1-β-D-ribofuranoside (DRB) and the translational inhibitor cyclohexamide (CHX) in combination with O/A for 24h in wild-type and PPM1H KO MEFs (Supplementary Figure 11, E). Consistent with the PPM1H mRNA time course, the O/A-induced increase in PPM1H protein levels was completely prevented when either transcription or translation were blocked (Figure 4 D, E). This was confirmed by RT-PCR analysis where the O/A-induced increase in PPM1H mRNA was abolished by transcription inhibition with DRB (Figure 4 F). Interestingly, translation inhibition led to a significant increase in basal PPM1H mRNA levels, and this increased further when mitochondria were depolarized following O/A treatment (Figure 4 F).

**Figure 4.**
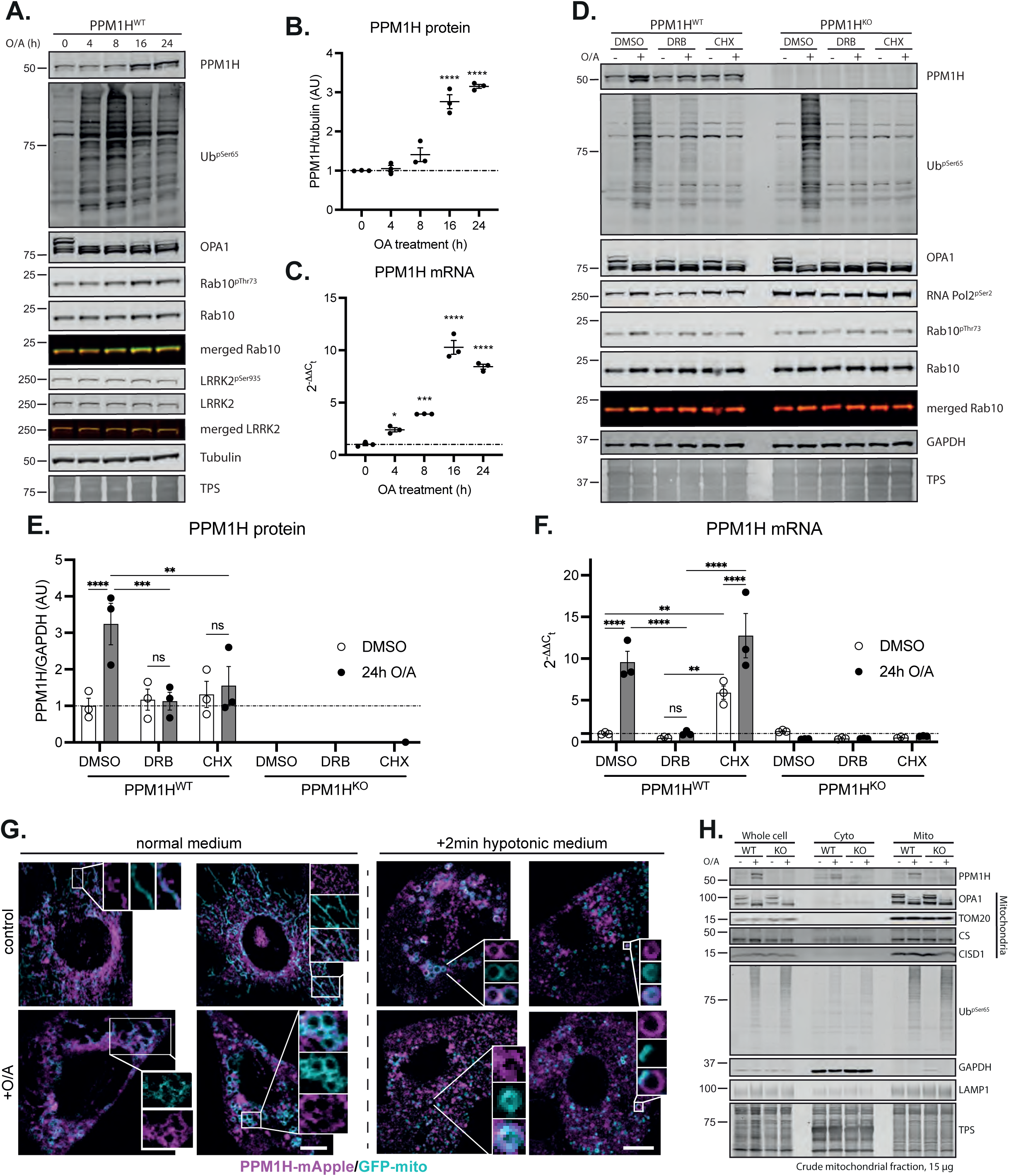
Endogenous PPM1H is up-regulated following mitochondrial depolarisation by a transcriptional mechanism. ***A.*** Representative immunoblot from PPM1H^WT^ MEFs treated with oligomycin/antimycin A for 0, 4, 8, 16, and 24 hours. ***B.*** Quantification of PPM1H protein and ***C.*** PPM1H mRNA from three independent experiments. ***D.*** Representative immunoblot from PPM1H^WT^ and PPM1H^KO^ MEFs upon 24h treatment with O/A, O/A+DRB or O/A+CHX. ***E.*** Quantification of PPM1H protein and ***F.*** PPM1H mRNA from three independent experiments. ***G.*** Live cell imaging of PPM1H-mApple MEF expressing a GFP mitochondrial tag treated with or without oligomycin/antimycin either in normal medium (left) or hypotonic buffer (right) ***H.*** Immunoblot analysis of whole cell extract, cytoplasmic fraction and crude mitochondrial fraction in PPM1H^WT^ and PPM1H^KO^ MEF after mitochondrial depolarization. Each lane was loaded with 40 μg (15 μg for mitochondrial fraction) of protein lysates. Graphs show mean ± SEM from three independent experiments. For ***B.*** and ***C.,*** ordinary one-way ANOVA with Dunnett’s multiple comparison test. For ***E.*** and ***F.,*** ordinary 2-way ANOVA with Tukey’s multiple comparison test from three independent experiments. Empty circles denote DMSO control while black circles represent O/A treated samples. Scale bar 50 μm.

PPM1H is predominantly localised at the Golgi with a small pool located at the mitochondria [16, 17]. Moreover, our prior analysis demonstrated that artificial localization of PPM1H to mitochondria blocks its ability to act on Thr72-phosphorylated Rab10 [17]. We next determined whether PPM1H is being targeted to mitochondria following O/A treatment. Live cell imaging studies of wild-type MEFs transiently co-transfected with PPM1H-mApple and GFP-mito revealed co-localisation on sites of fragmented mitochondria following O/A treatment (Figure 4 G). Furthermore, this co-localisation was readily detected upon 2 min treatment with hypotonic medium that facilitates identification of membrane contact sites (Figure 4 G). Immunoblotting analysis of mitochondrial fractions of MEFs confirmed basal expression of PPM1H and this increased following O/A treatment (Figure 4 H).

### Mutant PINK1 and LRRK2 exhibit convergent defects in ciliogenesis in the brain

We have previously reported that 7-month-old LRRK2 [R1441C] knock-in mice exhibit significantly fewer primary cilia in cholinergic interneurons within the dorsal striatum compared to wild-type littermate controls [11]. We therefore investigated whether endogenous PINK1 plays any role in the mutant LRRK2-mediated cilia defect and analysed ciliogenesis in dorsal striatal cholinergic interneurons from wild-type, LRRK2 [R1441C], PINK1 KO mice, and double mutant LRRK2 [R1441C] / PINK1 KO 5-month-old mice. We observed a small but significant loss of primary cilia in cholinergic interneurons of PINK1 KO mice, of a magnitude less than that seen in LRRK2 [R1441C] mice (Figure 5 A, C). Furthermore, we did not observe any exacerbation of the cilia loss in the double mutant LRRK2 [R1441C] / PINK1 KO mice, suggesting that the regulation of cilia by mutant LRRK2 and PINK1 occurs via parallel pathways (Figure 5 A, C). We next investigated ciliogenesis in striatal astrocytes where we have previously reported a ciliation defect in LRRK2 mutant models [55] and observed marked loss of cilia in astrocytes of PINK1 KO and LRRK2 [R1441C] mice (Figure 5 B, E) but again this was not worsened in the double mutant LRRK2 [R1441C] / PINK1 KO mice, suggesting that mutant LRRK2 and PINK1 exert parallel and convergent defects on cilia (Figure 5 B, E).

**Figure 5.**
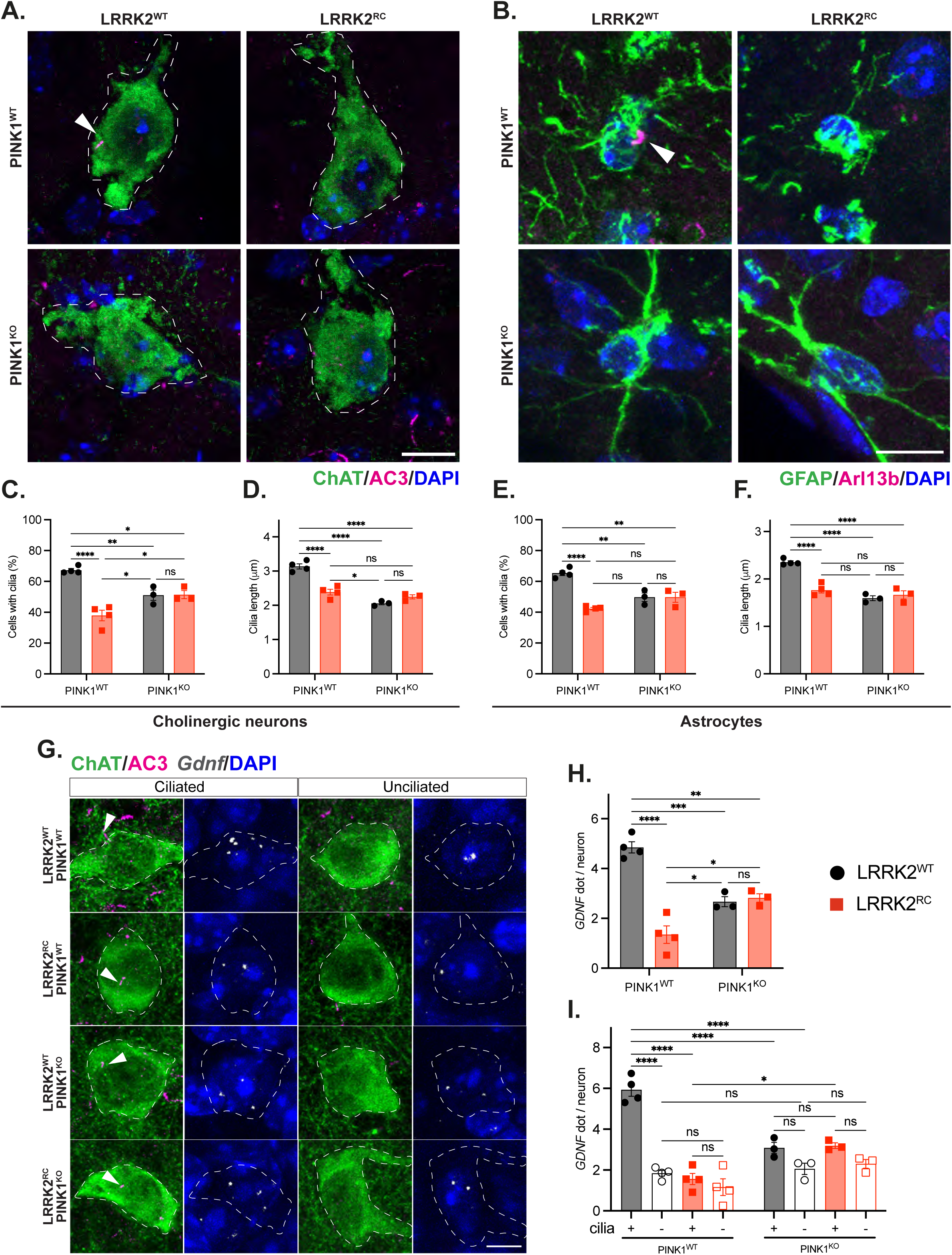
Loss of PINK1 decreases ciliary signalling in mouse striatum *in vivo*. ***A.*** and ***B.,*** Confocal images of sections of dorsal striatum from 5-month-old WT, LRRK2^RC^, PINK1^KO^ and LRRK2^RC^/PINK1^KO^ mouse brains. ***A***. Cholinergic interneurons were labelled using anti-choline acetyltransferase (ChAT) antibody (green), primary cilia were labelled using anti-AC3 antibody (magenta, white arrow), and nuclei were stained with DAPI (blue). ***B.*** Astrocytes were labelled using anti-GFAP antibody (green), primary cilia were labelled using anti-Arl13b antibody (magenta, white arrow) and nuclei were labelled using DAPI (blue). ***C.*** Quantitation of the percentage of ChAT^+^ neurons containing a cilium and ***D.*** their cilia length. Similar analyses for astrocytes are shown in ***E*.** and ***F. G.,*** Confocal images to identify ChAT^+^ neurons and their cilia as in ***A*** (left columns) coupled with RNAscope in situ hybridization to detect GDNF transcripts (right columns), segregated by ciliation status in WT, LRRK2^RC^, PINK1^KO^ and LRRK2^RC^/PINK1^KO^ mouse brains as indicated. ***H***. Quantitation of GDNF RNA dots per neuron or ***I.*** segregated as a function of ciliation status. Error bars represent SEM from N=3,4 mouse brains, with > 30 ChAT^+^ neurons and 25 astrocytes scored per brain. In bar charts, black circle represents LRRK2^WT^ while red squares LRRK2^RC^ mice. Ordinary 2-way ANOVA with Sidak’s multiple comparison test. * p<0.05, ** p<0.01, *** p<0.001, **** p<0.0001. Scale bar = 10µm.

Cilia shortening decreases ciliary signalling capacity. We found that PINK1 KO decreased cilia length 30% and this was not exacerbated by the additional presence of the LRRK2 [R1441C] mutation (Figure 5 D, F). We have shown that cilia are critical for Hedgehog signalling and production of GDNF by striatal cholinergic interneurons, providing neuroprotection for tyrosine hydroxylase-positive dopamine neurons [51]. In LRRK2 pathway mutant striatum, loss of cilia correlates with loss of Hedgehog-responsive gene expression, leading to decreased expression of Patched (PTCH1) and GDNF RNAs [51]. We therefore explored whether cilia loss influenced overall GDNF production. As shown in Figure 5 H, LRRK2 [R1441C] striatal cholinergic neurons showed a 5-fold decrease in GDNF RNA levels, as monitored by RNAscope fluorescence in situ hybridization (Fig. 5G). PINK1 KO cholinergic neurons showed a two fold decrease in GDNF expression, in either a wild type or LRRK2 [R1441C] background. When the data were further parsed according to ciliation status (Figure 5 G, I), GDNF expression correlated with the presence of a primary cilium in wild type cells, however even ciliated LRRK2 mutant neurons were defective in GDNF production. Ciliated PINK1 KO cells showed higher GDNF expression than unciliated PINK1 KO cells but again, even ciliated PINK1 KO cholinergic neurons displayed much less than wild type levels of GDNF expression. These findings are likely explained by the shorter cilia detected in these cholinergic neurons (Figure 5 D). These data demonstrate that PINK1 KO influences cholinergic ciliation and GDNF expression in the mouse dorsal striatum. Further work will be needed to explain why the PINK1 KO phenotype is not made more severe when combined with the LRRK2 [R1441C] mutation.

## DISCUSSION

Previous studies have established that LRRK2 lies within an endo-lysosomal signalling network with other PD gene-encoded proteins including VPS35, RAB29, RAB32 that act upstream of LRRK2 and in which disease-associated mutations lead to LRRK2 hyperactivation and increased Rab phosphorylation [14, 45, 58, 59]. Pathogenic activation of LRRK2 exerts downstream effects including lysosomal stress that confers cross-talk with additional PD-linked proteins including VPS13C and GBA1 [14, 60, 61] and this is associated with loss of primary cilia in selective cell types in the striatum of LRRK2 mutant mice [11, 55]. Endo-lysosomal pathways lie downstream of mitophagy and as outlined in the introduction there has been substantial interest in whether the LRRK2 pathway may interplay with the PINK1 pathway. Whilst our results show that knockout of endogenous PINK1 has no significant impact on endogenous LRRK2 activity *in vivo*, we observed striking ciliary defects and reduced GDNF signalling in the striatum of PINK1 KO mice brain that supports a convergent mechanism.

Based on *in vitro* studies of recombinant purified Rab8A protein, we had previously reported that PINK1-dependent phosphorylation of Ser111 at Rab8A leads to inhibition of LRRK2-mediated phosphorylation of Rab8A at Thr72 [36]. However, we were unable to confirm this interplay in primary MEFs under conditions of endogenous expression levels of Rab8A, PINK1 and LRRK2 (Supplementary Figure 9). This suggests that PINK1 and LRRK2 target different pools of Rab8A in cells, consistent with emerging data for their distinct localisations with LRRK2 recruitment to damaged lysosomes (or pericentriolar membranes) by Rab proteins, leading to enhanced phosphorylation of Rab8A (and Rab10) whereas PINK1 is recruited to sites of damaged mitochondria [27, 28, 34]. Compelling data for a role of LRRK2 on mitochondrial biology and thereby potential interplay with PINK1 is the demonstration that LRRK2 knockout mice have elevated basal mitophagy whilst LRRK2 [G2109S] knock-in mice have reduced mitophagy that can be rescued by LRRK2 inhibitors, in distinct central nervous system (CNS) cell types such as dopaminergic neurons [42, 43]. Further, Holzbauer demonstrated, in iPSC-derived neurons, that hyperactive LRRK2 mutations or PPM1H knock-out led to recruitment of the motor adaptor JIP4 to the autophagosomal membrane leading to abnormal activation of kinesin and disrupted transport that would inhibit axonal autophagy [62, 63]. Therefore, we cannot rule out interplay of the LRRK2 and PINK1 pathways in specific CNS cell types such as dopaminergic neurons. Furthermore, we cannot rule out that phosphorylation of alternate Rabs by LRRK2, not tested here, may be affected by PINK1. In future work it will be interesting to undertake a systematic analysis of Rab phosphorylation using unbiased proteomics approaches in select CNS cell types [47].

Both LRRK2 [R1441C] knock-in mice and PINK1 knock-out mice do not exhibit strong motor defects consistent with previous studies and critically the double mutant did not show any worsening of motor phenotypes (Figure 2). LRRK2 [R1441C] knock-in mice have been reported to be more susceptible to mitochondrial dysfunction [64]. Ultrastructural studies of mitochondria at the striatal pre-synaptic terminals of aged LRRK2 [R1441C] mice are reported to be abnormal with disrupted cristae and this is associated with reduced ATP production [64]. Interestingly, analysis of synaptic function in PINK1 KO and LRRK2 KO rats found age-dependent abnormalities in basal dopamine for both models and furthermore aged PINK1 rats showed significant disruption of neurotransmitter release with age-dependent increase in potassium evoked striatal dopamine release which was not observed in LRRK2 KO rats [65]. PINK1 KO mice have also been reported to exhibit abnormalities in neurotransmitter release [66] and in future work it would be interesting to determine whether there was any interplay between PINK1 and mutant LRRK2 in these synaptic defects.

Cholinergic interneurons are a rare subset of neurons in the striatum that sense and respond to Sonic Hedgehog secreted by dopaminergic neurons; in turn, these cells secrete GDNF to provide trophic support for dopaminergic neurons [67]. Previous work has revealed that hyperactive mutants of LRRK2 including R1441C and G2019S lead to loss of primary cilia in cholinergic interneurons and that this can be detected at 10 weeks of age [11, 55]. Furthermore, loss of the Rab phosphatase, PPM1H, exhibits a similar ciliary defect providing strong genetic evidence for an important role for LRRK2 pathway activity in cilia formation [55]. We report for the first time that loss of PINK1 can also lead to primary ciliary loss in striatal cholinergic interneurons and astrocytes, however, we did not observe an exacerbation of the ciliary loss in the double mutant LRRK2 R1441C/PINK1 KO mice (Figure 5). Moreover, loss of PINK1 also led to ciliary shortening, the consequence of which led to significantly decreased Hedgehog signalling and decreased GDNF RNA production. These findings imply parallel routes to a convergent pathway between LRRK2 mutations and PINK1 knockout, both triggering loss of neuroprotection in the dorsal striatum by independent routes.

The mechanism of how the PINK1 pathway impacts on cilia is unclear. It was recently reported that human iPSC-derived neuronal precursor cells PINK1 KO mice striatal neurons exhibit shortened primary cilia defects [68]. Furthermore, it has been reported that mitochondrial stress, that can be induced by inhibitors of mitochondrial respiration chain complexes, can stimulate ciliogenesis in a variety of CNS cell types mediated via reactive oxygen species [69]. It has also been reported that mtDNA loss in astrocytes lacking the Twinkle helicase exhibit abnormal, elongated and more motile cilia associated with mitochondrial respiratory chain deficiency and aberrant transcription [70]. In future work it will be interesting to investigate the mechanism of cilia regulation PINK1.

In prior work we found that endogenous PPM1H levels were increased in mitochondria of primary mouse cortical neurons following mitochondrial depolarisation induced by O/A treatment [50]. By immunoblotting, we observed an increase in PPM1H levels in MEF whole cell extracts following O/A treatment and subcellular fractionation and live cell imaging studies revealed higher recruitment of PPM1H to the mitochondrial membrane following O/A treatment. Furthermore, the PPM1H response to mitochondrial depolarisation was independent of PINK1 and LRRK2 catalytic activity. Over-expression studies have previously revealed that PPM1H is localised mainly to the Golgi complex with further pools of PPM1H associated with the mother centriole and mitochondria although PPM1H does not act on mitochondrial Ser111-phosphorylated Rab8 [16, 17]. At the Golgi, PPM1H strongly co-localises with Rab8A, Rab10 and Rab29 but less well with Rab12 and over-expression of PPM1H efficiently dephosphorylates Rab10 but not Rab12 suggesting that its major role is to protects Golgi-associated Rabs from LRRK2 phosphorylation and inactivation [17]. Herein, we observed that the increase in PPM1H expression by mitochondrial depolarisation was not accompanied by a concomitant reduction in phosphorylated Rab10 and this is in line with a previous study in which artificial tethering of PPM1H to the mitochondria led to impaired ability to dephosphorylate total Rab10 [17]. In future studies it will be interesting to better understand the functional consequence of stress-induced PPM1H recruitment to the mitochondria and whether this is important for mitigating LRRK2-dependent phosphorylation of yet unidentified Rabs at the mitochondrial membrane as part of a protective response. Recently autosomal dominant mutations in Rab32 have been identified as a cause of PD and it has further been shown that Rab32 mutations lead to LRRK2 activation and Rab phosphorylation [59]. Previous studies have indicated that Rab32 is located at the mitochondria [71–73] and a recent study has demonstrated that LRRK2 forms a complex with

Rab32 and aconitate decarboxylase 1 (IRG1) at the mitochondria that is enhanced by Salmonella infection and this complex is critical for delivery of antibacterial aconitase from the mitochondria to salmonella containing vesicles [74]. There are common mechanisms by which cells respond to mitochondrial stress and bacterial pathogen infection e.g. clearance of damaged mitochondria by mitophagy or bacteria by xenophagy [75]. In future work it would be exciting to investigate the potential role of PPM1H at stressed mitochondria and whether this mitigates mutant Rab32 mediated LRRK2 substrate phosphorylation. Further, it would be interesting to determine whether PPM1H is up-regulated in response to Salmonella infection to counteract the protective role of the LRRK2-Rab32-IRG1 complex.

In summary, endogenous LRRK2 and PINK1 function in parallel signalling pathways *in vivo*, however, mutation of both genes leads to impaired ciliogenesis in the brain suggesting a convergent neurobiological mechanism for Parkinson’s disease gene pathways. There is growing interest in delivering GDNF to PD patients as a therapeutic strategy and our findings would suggest that both PINK1 and LRRK2 mutant carriers may benefit from such targeted therapies. In contrast, there are several clinical trials underway for evaluating whether LRRK2 inhibitors or antisense oligonucleotide therapies (ASOs) confer disease-modifying benefits for PD patients [76, 77] and our analysis would suggest that patients harbouring PINK1 mutations would not benefit from LRRK2 inhibitors and highlight the need for patient stratification for molecular targeted clinical trials in PD.

## MATERIALS AND METHODS

All antibodies, chemicals and reagents, and mouse strains are listed separately in Supplementary Key Reagents Table.

### Animal husbandry

Mice were housed in temperature-controlled rooms at 21°C with 45-65% relative humidity, 12h/12h light/dark cycle and ad libitum access to food and water. All mice in this study had automatic watering (0.2 micron sterile filtered) and were fed rodent diet ‘‘R&M No.3, 9.5 mm pelleted (irradiated, Special Diets Services, UK). All cages had corn-cob substrate (provided as a nest-pack) and sizzle nest material, additionally environmental enrichment was provided for all animals, with a cardboard tunnel for amalgamated females, single-housed males and squabbling males. Cages were changed as needed, but all cages were changed on at least a two-weekly cycle while mice were regularly subjected to health and welfare monitoring as standard (twice-daily). All mice in this study were maintained on a C57BL/6J background. Mice of both genders were used in all experiments. All animal studies were approved by the University of Dundee Ethical Review Committee and performed under a U.K. Home Officer project license. Experiments were conducted in accordance with the Animal Scientific Procedures Act (1986) and with the Directive 2010/63/EU of the European Parliament and of the Council on the protection of animals used for scientific purposes (2010, no. 63).

### Mice behavioural and motor test

Behavioural tests were conducted on 10.5 months old mice. Mice were weighted before the start of behavioural tests to make a comparison between genotypes.

Negative geotaxis was assessed by placing the animal onto a mesh grid (30 x 30 cm). The time taken to rotate through 180° from a head down position was recorded as a measure of proprioception.

Grip strength was measured using a grip meter modified from GSM1054 model (Linton Instrumentation) as previously described [78]. In two consecutive trials, the mouse was held by the tail and lowered onto the instrument until it gripped the two bars. The mouse was pulled by the base of the tail until the grip loosened. The applied force at which the mouse released the bars was recorded and averaged across the two trials.

Gait analysis, rotarod and balance beam were conducted as described in [47]. Briefly, gait analysis was carried out using the footprint test. The animal was placed in a clear Perspex corridor apparatus (65 cm L x 15 cm W) and trained to run towards a dark goal box at the end of the corridor until it could reach the box without encouragement. For testing, a paper strip was placed in the corridor and, to leave footprints, the mouse’s paws were painted with non-toxic, water-based paints in two different colours to identify the front paws versus the hind paws. The mouse was allowed to run the entire length of the apparatus and reach the goal box. The stride length, the stride width and the overlap were measured using 4 paws print, allowing to average 3 values for each measurement.

Rotarod was carried out using a commercial Rotarod apparatus (Ugo Basile, model 47600). After 5 sessions of training conducted over 5 consecutive days (max 5 min/session), mice were tested in two different trials (accelerating rod from 5 rpm to 44 rpm in each trial). The latency to fall was recorded and averaged across the two trials. For the balance beam, mice were trained on an elevated bridge (1 m in length, 17% angle of ascent, with 1.5 to 0.5 cm tapers across the width) with a dark house box at the high end. During the first day of training, the mouse was placed in front of the house box and allowed to enter the box. The distance from the house box was progressively increased until the low end of the beam. The mouse was then placed at the low end of the beam, facing away from the house box, and encouraged to turn around and transverse the beam until the house box. The test was carried out in two consecutive trials, conducted 1 hour apart, and videotaped to allow analysis. The mouse was placed at the low end of the beam, facing away the house box. The time taken to turn around, transverse the beam and the number of foot slips were recorded and averaged across the two trials.

### MLi-2 treatment in mice

To ensure LRRK2-dependent phosphorylation of Rabs, mice were treated with the LRRK2 inhibitor MLi-2. The compound was administered to mice via subcutaneous injection as described (dx.doi.org/10.17504/protocols.io.bezdjf26). MLi-2 was resuspended in a 40% Hydroxypropyl-β-Cyclodextran (Average Mw ∼1,460) solution at 6 mg/ml. It was then administered by subcutaneous injection at 30 mg/kg. The Dundee-synthesised MLi-2 (MTA-free) was used for this experiment. Mice were culled 2 hours after the injections, tissue collected and lysed as outlined above.

### Mouse brain immunohistochemistry – fluorescence analysis

Analysis of primary cilia in the mouse brain striatum was performed as previously described (dx.doi.org/10.17504/protocols.io.bnwimfce). Mice were anaesthetised using a commercial solution of Euthatal, before being perfused with PBS and 4% PFA. The brain was then dissected, fixed overnight in 4% PFA at 4°C, washed and left in 30% sucrose for 48h at 4°C. Whole brains were subsequently embedded in 22 x 22 x 20 mm molds containing O.C.T. compound and kept at -80°C until sectioning. Sections of the mouse striatum were then obtained with a cryostat with a cutting thickness of 16 µm. Frozen sections were thawed at RT for 15 min and gently washed (2X) with PBS for 5 min. For antigen retrieval, slides were incubated with 10 mM sodium citrate buffer pH 6.0 (preheated to 95° C) for 15 minutes at 95° C. Sections were permeabilized with 0.1% Triton X-100 in 1X PBS at RT for 15 min. Sections were blocked with 2% FBS and 1% BSA in PBS for 2 hr at RT and were then incubated overnight at 4°C with primary antibodies. The following day, sections were incubated with secondary antibodies at RT for 2 hr. Donkey highly cross-absorbed H + L secondary antibodies conjugated to Alexa 488 and Alexa 568 were used at a 1:2000 dilution. Nuclei were stained with 0.1 µg/ml DAPI (Sigma). Stained tissues were overlaid with Fluoromount G and a glass coverslip. All antibody dilutions for tissue staining included 1% DMSO to help antibody penetration. Images were obtained using a Zeiss LSM 900 confocal microscope with a 63 x 1.4 oil immersion objective. Image visualisations and analyses were performed using Fiji.

### Mouse brain immunohistochemistry - colorimetric analysis

For Iba1 and DARPP-32 staining, mice were anesthetised with Euthatal and perfused with PBS and PFA 4%. Brains were fixed overnight in 4% PFA at 4°C, washed and left in 30% sucrose for 24h at 4°C. Brains were sliced into 35 µm-thick slices using a freezing microtome and stored at -20°C until processing for immunohistochemistry. Free-floating sections were rinsed (3X) with TBS for 10 min and incubated with quenching solution (3% H_2_O_2_, 10% Methanol in TBS) for 15 min. Sections were subsequently rinsed (3X) in TBS for 10 min and incubated with blocking solution (5% normal goat serum, TBS-Triton 0.1%) for 1 h at room temperature. Incubation with primary antibodies was performed overnight at 4°C. The following day, sections were rinsed (3X) with TBS Triton 0.1% for 10 min and incubated with the secondary antibodies for 2 h at room temperature. Sections were subsequently rinsed (3X) in TBS-Triton 0.1% for 10 min and incubated for 2 h at room temperature with avidin-peroxidase complex (ABC kit, PK4000, Vector). 3,3′-diaminobenzidine (DAB, Sigma) was applied to the slices to visualise Iba1 and DARPP-32 positive cells. Imaged were obtained using a bright-field microscope (Macro/Micro Imaging System, Leica) under a 40X objective and analysed using Fiji.

### Fluorescence in situ hybridization (FISH)

RNAscope fluorescence in situ hybridization was conducted as described herein: (bio-protocol.org/prep1423) [51, 55]. The RNAscope Multiplex Fluorescent Detection Kit v2 (#323100, Advanced Cell Diagnostics) was used as per the manufacturer with RNAscope 3-plex Negative Control Probe (#320871) or probe Mm-Gdnf-C1 (#421951). The Mm-Gdnf-C1 probe was diluted 20 X in a buffer containing 6x saline-sodium citrate (SSC), 0.2% lithium dodecyl sulfate, and 20% Calbiochem OmniPur Formamide. Fluorescent visualization of the hybridized probes was achieved using Opal 690 (Akoya Biosciences). Brain slices were blocked with 1% BSA and 2% FBS in Tris-buffered saline with 0.1% Triton X-100 for 30 min. They were then incubated overnight at 4°C with primary antibodies in TBS containing 1% BSA and 1% DMSO. This was followed by treatment with secondary antibodies, diluted in TBS with 1% BSA and 1% DMSO, including 0.1 µg/ml DAPI (Sigma) for 2h at room temperature. Finally, sections were mounted using Fluoromount G and glass coverslips.

### Cell and tissue lysis and immunoblotting

Cells were quickly washed on ice in PBS, then lysed in buffer containing Tris-HCl (50 mm, pH 7.5), EDTA (1 mM), EGTA (1 mM), Triton (1% w/v), sodium orthovanadate (1 mM), sodium glycerophosphate (10 mM), sodium fluoride (50 mM), sodium pyrophosphate (10 mM), sucrose (0.25 mM), protease inhibitor cocktail (Roche), phoSTOP (Roche), and chloroacetamide (200 mM). Tissues were instead collected and snap frozen in liquid nitrogen. They were then weighted, quickly thawed on ice in a 10-fold volume excess of ice cold lysis buffer. Tissues were homogenised using a POLYTRON homogenizer (KINEMATICA), employing three rounds of 10s homogenization with 10s intervals on ice. Lysates either from cells or tissues, were incubated for 30 min on ice. Samples were spun at 17000 g in an Eppendorf 5417R centrifuge for 30 min at 4°C. Supernatants were collected, and protein concentration was determined by using the Bradford kit (Pierce).

Protein lysates were subjected to SDS–PAGE (4–12% Bis-Tris gel or 12% Tris glycine) and transferred onto nitrocellulose membranes. Membranes were then blocked for 1 h in Tris-buffered saline with 0.1% Tween (TBST) containing 5% (w/v) milk and subsequently probed with the indicated antibodies in TBST containing 5% (w/v) BSA overnight at 4°C. Detection was performed using appropriate secondary antibodies (1:10000) and scanned using Li-COR Odyssey CLx imaging system. More details can be found on protocols.io (dx.doi.org/10.17504/protocols.io.ewov14znkvr2/v2). Signal intensity was quantified using the Image Studio Software and normalised versus the unphosphorylated protein or the loading control. The amount of protein loaded in each lane is reported for each blot.

### pSer65 Ub ELISA

Phosphorylation of Ub at Ser65 by PINK1 was monitored *in vivo* by enzyme linked immunosorbent assay (ELISA) as previously described by Watzlawik and colleagues [54]. MSD plates were coated overnight with 30 μl/well of 200 mM sodium carbonate buffer (pH 9.7) containing 1 μg/ml of rabbit monoclonal pSer65-Ub antibody. The next morning plates were washed twice with ELISA washing buffer (150 mM Tris, pH 7.4, 150 mM NaCl, 0.1% Tween-20) by plate inversion and gentle tapping on paper towels (not by pipette aspiration). Plates were then blocked with ELISA blocking buffer (150 mM Tris, pH 7.4, 150 mM NaCl,0.1% Tween-20, 1% BSA) for 1h at room temperature. All samples were run in duplicates and diluted in blocking buffer. 30 μg of total protein were loaded per well for all mouse tissues in a total volume of 30 μl per well. Detergent volumes were adjusted across all samples. Antigens were incubated for 2 h at room temperature on a microplate mixer at 600 rpm and three washing steps were then performed as described before. Mouse total Ub antibody (clone P4D1; Thermo Fisher #14-6078-37) was subsequently added as detecting antibody at a final concentration of 5 µg/ml in blocking buffer in 30 μl total volume per well. After three washing steps, 50 μl/well of 1 μg/ml of SULFO-TAG labelled goat anti-mouse antibody (MSD, R32AC-1) in blocking buffer were added and incubated for 1 h at room temperature on a microplate mixer at 500 rpm. After another three washing steps, 150 μl MSD GOLD Read Buffer (MSD, R92TG-2) were finally added to each well and the plate read on a MESO QuickPlex SQ 120 reader.

### MEF generation, maintenance, and treatment

Mouse embryonic fibroblasts (MEFs) were isolated from mice embryos of different genotype and relative littermate WT as extensively described in dx.doi.org/10.17504/protocols.io.eq2ly713qlx9/v1 and then immortalised by SV40-mediated immortalization. MEFs were cultured in DMEM high glucose supplemented with 10% (v/v) FBS, 2 mM L-glutamine, Penicillin-Streptomycin 100U/mL, 1 mM Sodium Pyruvate, 1X NEAA solution. To induce mitochondrial depolarization, MEFs were treated for 24h with a combination of Oligomycin and Antimycin A at a final concentration of 1 μM and 10 μM. The type-2 LRRK2 inhibitor MLi-2 was used at a final concentration of 100 nM for 1h and 30 min. Inhibition of transcription and translation was achieved by using 5,6-dichlorobenzimidazole (DRB) at 100 μM and cycloheximide at 1 μg/ml for 24h. All compounds were dissolved and diluted in DMSO.

### PINK1 siRNA

PINK1 knockdown was performed by siRNA in 6 well plates as extensively described (at DOI: dx.doi.org/10.17504/protocols.io.kxygx343zg8j/v1). Briefly, 100000 cells were seeded in each well for both LRRK2^WT^ and LRRK2^RC^ MEFs. The following day, cells were either incubated with 25 nM of either mouse PINK1 siRNA or scrambling siRNA (Dharmacon). After 48 hours, oligomycin and antimycin were added to the culture medium at a final concentration of 1 μM and 10 μM and incubated for another 24 hours. The next day (4 days after cell seeding), the LRRK2 kinase inhibitor MLi-2 was added at a concentration of 100 nM for 1h and 30 min. Finally, cells were quickly washed and lysed on ice, the protein concentration quantified by Bradford and the lysates subjected to western blotting.

### Mitochondrial fractionation

Mitochondrial fractions were purified following steps 21-32 of dx.doi.org/10.17504/protocols.io.bxmypk7w. Briefly, two 15 cm^2^ dishes (for each sample) were scraped on ice and collected in hypotonic buffer [20 mM Hepes (pH 7.8), 5 mM potassium chloride, 1.5 mM magnesium chloride, 2 mM DTT, 1 mM PMSF and both protease and phosphatase inhibitor cocktails (Roche)]. Cells were homogenised with 45 strokes of stainless steel dounce homogeniser, then, 2.5× mannitol-sucrose buffer [2.5× MSH; 20 mM Hepes (pH 7.8), 525 mM mannitol, 175 mM sucrose, 5 mM EDTA, 1 mM PMSF, and protease inhibitor cocktail (Roche)] was added to the disrupted cells, and the cell homogenates were clarified by centrifugation (700g at 4°C for 10 min) to remove nuclei and cell debris. Supernatants were collected and spun down again at 700g at 4°C for 10 min before mitochondria were pelleted at 9000g for 10 min. The pellet was then resuspended and washed twice in 1× MSH [20 mM Hepes (pH 7.8), 210 mM mannitol, 70 mM sucrose, 2 mM EDTA, 1 mM PMSF, and protease inhibitor cocktail (Roche)] and centrifuged at 9000g for 10 min at 4°C. Finally, mitochondrial pellets were resuspended in 50 μl of lysis buffer, protein quantified, and lysates interrogated by western blotting.

### Live cell imaging

Intracellular localization of PPM1H was monitored in MEFs stably expressing a fluorescent PPM1H-mApple and a GFP-tagged to monoamine oxidase A (MAO-A). Cells were seeded in 8-well glass incubation chambers (5000 cells/well, Nunc 155409) with 0.2 ml of cultured medium. The following day, culture medium was exchanged for phenol red-free medium, with or without oligomycin and antimycin (4hr) and/or hypotonic buffer consisting of 5% DMEM in sterile H_2_O (2 min). The 8-well chamber was then placed onto a heated microscopy stage with CO_2_ supply and images were taken using confocal z-sectioning.

### RT-PCR

PPM1H mRNA were quantified by RT-PCR as previously described (dx.doi.org/10.17504/protocols.io.81wgbz7r3gpk/v1). Briefly, 15cm^2^ dishes were washed twice in DPBS and cells scraped in 1ml of DPBS. Each sample was then divided, 2/3 were used for western blots and lysed as previously described. The remaining third was spun down, the supernatant removed, and the cell pellet snap frozen and used for RT-PCR. RNA was extracted using the PureLink™ RNA Mini Kit (ThermoFisher) and following manufacturer’s instructions. Cells were dissociated using 300 μl of lysis buffer and by 10x passages in a 20G needle. Total RNA was then eluted in 50 μl of molecular biology water and stored at -80°C degree. cDNA synthesis was achieved using the QuantiTect Reverse Transcription Kit (Qiagen) using 1 μg of RNA as template and following manufacturer’s instructions. The obtained cDNA was diluted 1 in 5 before used for RT-PCR. Two pairs of primers were used to assess PPM1H, while 4 housekeeping genes were used for normalization (ACTB, GAPDH, RPL13A and TBP). In each well of a 384 well plate, 2 μl of cDNA were mixed with 3 μl of PowerUp™ SYBR™ Green Master Mix (ThermoFisher) containing 1 μM of forward and reverse primer. The plate was then placed in a thermocycler, the C_t_ value extrapolated from the amplification curves and the data analysed using the ΔΔC_t_ method.

### Statistical analysis

Data were analysed and plotted using Prism 10.0.3. Statistical difference, set at pL<0.05, was calculated either by ordinary one-way ANOVA or by ordinary two-way ANOVA with the appropriate multiple correction test. For not-normally distributed data, a non-parametric test was used instead. P values ≤ 0.05, 0.01, 0.001, 0.0001 are represented as *, **, ***, **** respectively. Graphs represent mean ± SEM, unless otherwise stated. The details of each statistical test, n numbers and graph used are reported in the relative figure legends.

## Supporting information

Supplementary Key Reagents Table

## ACKNOWLEDGEMENTS

We thank the technical support of the MRC Protein Phosphorylation and Ubiquitylation Unit (PPU) Genotyping team (coordinated by G. Gilmour and latterly Fiona Brown), the MRC-PPU DNA sequencing service (coordinated by G. Hunter), the MRC-PPU tissue culture team (coordinated by E. Allen), the MRC-PPU MS facility team (coordinated by R. Soares), and the MRC-PPU Reagents and Services team (coordinated by J. Hastie). This research was funded by Aligning Science Across Parkinson’s (ASAP-000463) through the Michael J. Fox Foundation for Parkinson’s Research (MJFF) (to M.M.K.M., D.R.A., S.R.P.), the Medical Research Council (MRC) MR/P00704X/1 COEN award (to M.M.K.M., D.R.A.), the pharmaceutical companies supporting the Division of Signal Transduction Therapy Unit (Boehringer Ingelheim, GlaxoSmithKline, and Merck KGaA (M.M.K.M. & D.R.A), and a Wellcome Trust Senior Research Fellowship in Clinical Science (210753/Z/18/Z to M.M.K.M.). The M.M.K.M. laboratory is supported by the Michael J. Fox Foundation and an EMBO YIP Award. The D.R.A. laboratory is also supported by the UK Medical Research Council (grant number MC_UU_00018/1). R.B. laboratory is supported by the Medical Research Council Award (MR/S037667/1) of the “Therapeutic Target Validation in Mental Health” Programme. Schematics for figures were created using Biorender.

## ABBREVIATIONS

Arl13b: ADP ribosylation factor like GTPase 13B
CHX: Cycloheximide
DRB: 5,6-Dichlorobenzimidazole 1-β-D-ribofuranoside
GDNF: glial-derived neurotrophic factor
GFAP: Glial fibrillary acidic protein
KO: Knock-out
LRRK2: Leucine-rich repeat kinase 2
MEFs: Mouse embryonic fibroblasts
O/A: Oligomycin/Antimycin A
PD: Parkinson’s disease
PINK1: PTEN-induced kinase 1
pSer: Phospho-serine
pThr: Phospho-threonine
RC: R1441C
TPS: Total protein stain
Ub: Ubiquitin

## SUPPLEMENTARY FIGURE LEGENDS

**Supp Figure 1**

**LRRK2 signalling in the olfactory bulb and hippocampus is not affected by loss of PINK1 *in vivo. A.*** Immunoblot of LRRK2 pathway component in mouse olfactory bulb and ***B.*** hippocampus. Quantification of ***C***. pSer105/total Rab12, ***D***. PPM1H/Vinculin and ***E.*** pSer935/total LRRK2 for the olfactory bulb and in ***F., G., H.*** for the hippocampus. Each lane was loaded with 40 μg of protein lysate from one mouse. In graphs, black circle represents PINK1^WT^ while red square PINK1^KO^ animals. Box and whiskers plot, from min to max with the median line. Ordinary 2-way ANOVA with Sidak’s multiple comparison test. * p<0.05, ** p<0.01, *** p<0.001, **** p<0.0001.

**Supp Figure 2**

**LRRK2 signalling in the midbrain and thalamus is not affected by loss of PINK1 *in vivo. A.*** Immunoblot of LRRK2 pathway component in mouse midbrain and ***B.*** thalamus. Quantification of ***C***. pSer105/total Rab12, ***D***. PPM1H/Vinculin and ***E.*** pSer935/total LRRK2 for the midbrain and in ***F., G., H.*** for the thalamus. Each lane was loaded with 40 μg of protein lysate from one mouse. In graphs, black circle represents PINK1^WT^ while red square PINK1^KO^ animals. Box and whiskers plot, from min to max with the median line. Ordinary 2-way ANOVA with Sidak’s multiple comparison test. * p<0.05, ** p<0.01, *** p<0.001, **** p<0.0001.

**Supp Figure 3**

**LRRK2 signalling in the cerebellum and brainstem is not affected by loss of PINK1 *in vivo. A.*** Immunoblot of LRRK2 pathway component in mouse cerebellum and ***B.*** brainstem. Quantification of ***C***. pSer105/total Rab12, ***D***. PPM1H/Vinculin and ***E.*** pSer935/total LRRK2 for the cerebellum and in ***F., G., H.*** for the brainstem. Each lane was loaded with 40 μg of protein lysate from one mouse. In graphs, black circle represents PINK1^WT^ while red square PINK1^KO^ animals. Box and whiskers plot, from min to max with the median line. Ordinary 2-way ANOVA with Sidak’s multiple comparison test. * p<0.05, ** p<0.01, *** p<0.001, **** p<0.0001.

**Supp Figure 4**

**LRRK2 signalling in the spinal cord is not affected by loss of PINK1 *in vivo. A.*** Immunoblot of LRRK2 pathway component in mouse spinal cord. Quantification of ***B.*** pSer105/total Rab12, ***C.*** PPM1H/Vinculin and ***D.*** pSer935/total LRRK2. Each lane was loaded with 40 μg of protein lysate from one mouse. In graphs, black circle represents PINK1^WT^ while red square PINK1^KO^ animals. Box and whiskers plot, from min to max with the median line. Ordinary 2-way ANOVA with Sidak’s multiple comparison test. * p<0.05, ** p<0.01, *** p<0.001, **** p<0.0001.

**Supp Figure 5**

**LRRK2 signalling in the lung and spleen is not affected by loss of PINK1 *in vivo. A***. Immunoblot of LRRK2 pathway component in mouse lungs and relative quantification of ***B.*** pThr73/total Rab10, ***C.*** pSer105/total Rab12, ***D.*** pSer935/total LRRK2, and ***F.*** PPM1H/Vinculin. Similarly in ***F., G., H.***, and ***I.*** analysis from mouse spleen. pSer105 Rab12 was undetectable in this tissue. Each lane was loaded with 40 μg of proteins lysate from one mouse. In graphs, black circle represents PINK1^WT^ while red square PINK1^KO^ animals. Box and whiskers plot, from min to max with the median line. Ordinary 2-way ANOVA with Sidak’s multiple comparison test. * p<0.05, ** p<0.01, *** p<0.001, **** p<0.0001.

**Supp Figure 6**

Behavioural testing sub-analysis in 10.5 months old double mutant PINK1 knockout / LRRK2 R1441C mutant mice does not suggest genetic interaction *in vivo*. *A.* Forelimb and *B.* hindlimb paws base width during gait analysis. The overlap of the two is shown in graph *C*. The average amount of footslips after two balance beam tests is shown in *D.* for forelimbs. *E.* for hindlimb and their sum in *F.* Weight at 10.5 months separated by sex of the animal for females *G.* and males *H.* In violin plots, black circle represents LRRK2^WT^ while red squares LRRK2^RC^ mice. Ordinary 2-way ANOVA with Sidak’s multiple comparison test. * p<0.05, ** p<0.01, *** p<0.001, **** p<0.0001. N=15/16 mice per group

**Supp Figure 7**

**Immunohistochemistry analysis of medium spiny neurons in double mutant PINK1 knockout / LRRK2 R1441C mutant mice. *A.*** Representative images and ***B.*** quantification of DARPP32 staining and ***C.*** striatal volume in the brain of 10.5 months old mice. In bar charts, black circle represents LRRK2^WT^ while red squares LRRK2^RC^ mice. Ordinary 2-way ANOVA with Sidak’s multiple comparison test. N=15/16 mice per group. Scale bar 50 μm.

**Supp Figure 8**

**pSer65 Ubiquitin analysis in double mutant PINK1 knockout / LRRK2 R1441C mutant mice.** Detection of pSer65 Ub by ELISA in different regions of the CNS such as cortex ***A***., midbrain ***B.***, cerebellum ***C.*** and spinal cord ***D.*** In box and whiskers plots, black circle represents LRRK2^WT^ while red square LRRK2^RC^ mice. Ordinary 2-way ANOVA with Tukey’s multiple comparison test. N=4 mice per group.

**Supp Figure 9**

Analysis of LRRK2 and PINK1 Rab phosphorylation in primary MEFs derived from double mutant PINK1 knockout / LRRK2 R1441C mutant mice. *A.* Immunoblot of two independent MEF clones from WT/WT, KO/WT, WT/RC and KO/RC mice upon 24h treatment with O/A, 2h treatment with MLi-2. *B.* Quantification of pSer111 and *C.* pThr72 Rab8A following Rab8A immunoprecipitation. *D.* Quantification of pThr73 Rab10 and *E.* pSer935 LRRK2. In graphs, black circles represent LRRK2^WT^ while red square LRRK2^RC^ samples.

**Supp Figure10**

**Mitochondrial depolarisation-induced PPM1H increase is independent of PINK1 and LRRK2 in MEFs**. Immunoblot from ***A.*** PPM1H^WT^ and PPM1H^KO^, ***B.*** LRRK2^WT^ and LRRK2^KO^ and ***C.*** PINK1^WT^ and PINK1^KO^ MEFs upon 24h treatment with O/A, 2h treatment with MLi-2 or the combination of both. Each lane was loaded with 40 μg of protein lysates.

