## Supplementary Figures for "Endogenous LRRK2 and PINK1 function in a convergent neuroprotective ciliogenesis pathway in the brain"

#### Supp1

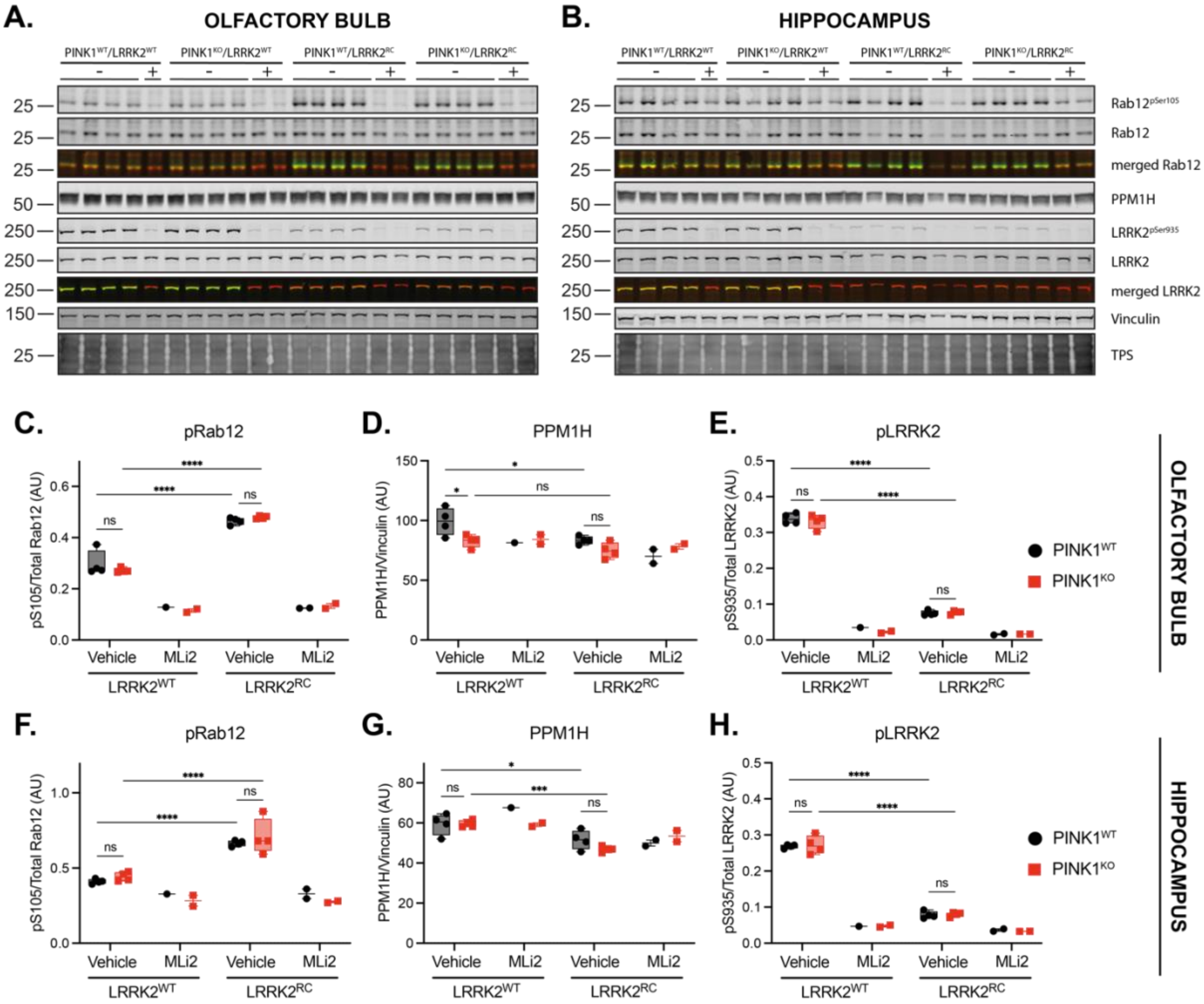

Supp2

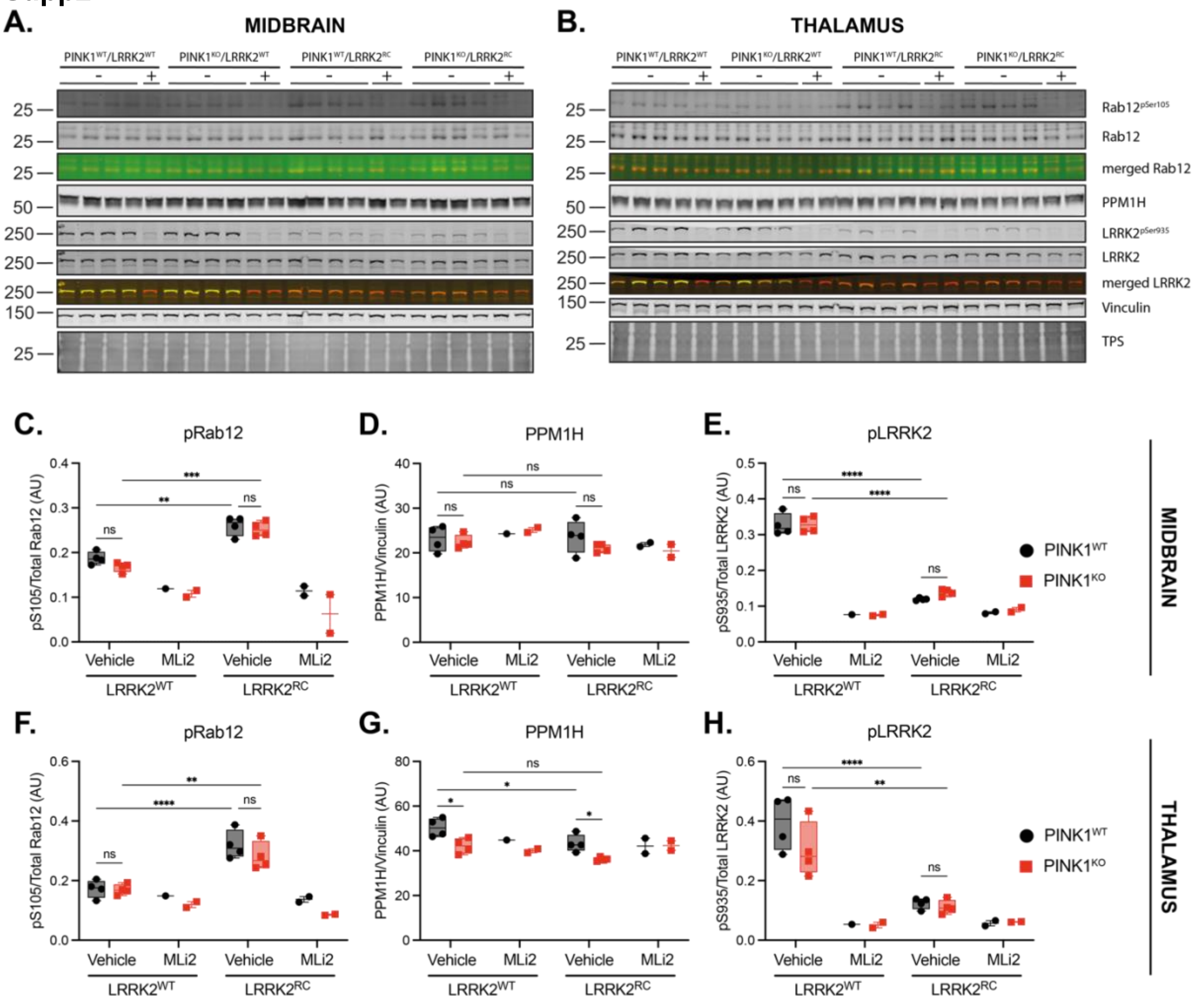
A.
MIDBRAIN
B.
THALAMUS
PINK1WT/LRRK2WT
PINK1KO/LRRK2WT
PINK1WT/LRRK2RC
PINK1KO/LRRK2RC
Rab12pSer105
Rab12
merged Rab12
PPM1H
LRRK2pSer935
LRRK2
merged LRRK2
Vinculin
TPS
C.
pRab12
D.
PPM1H
E.
pLRRK2
F.
pRab12
G.
PPM1H
H.
pLRRK2
pS105/Total Rab12 (AU)
PPM1H/Vinculin (AU)
pS935/Total LRRK2 (AU)
Vehicle
MLi2
LRRK2WT
LRRK2RC
PINK1WT
PINK1KO
MIDBRAIN
THALAMUS

**Supp3**

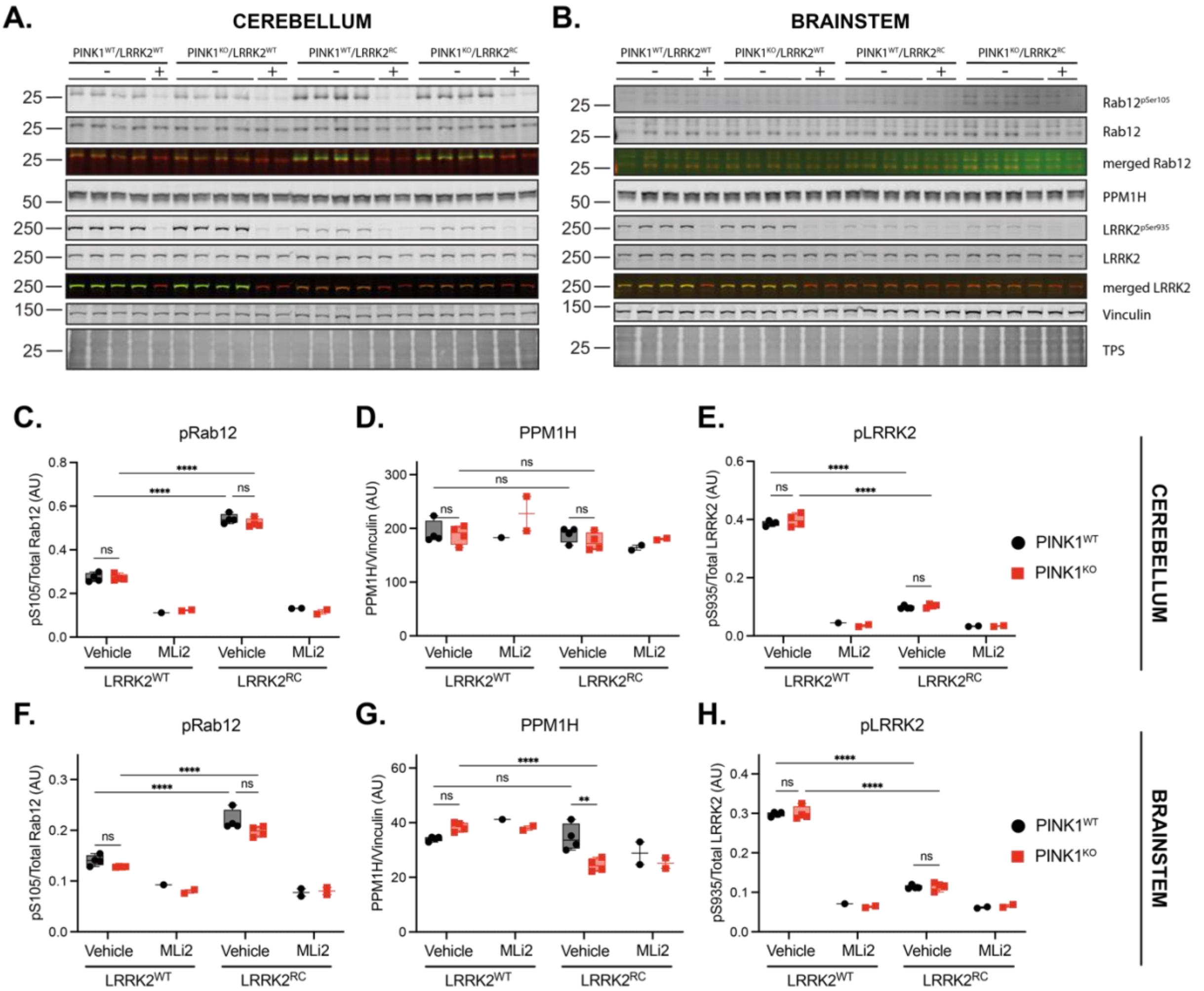

A.
CEREBELLUM
B.
BRAINSTEM
PINK1WT/LRRK2WT
PINK1KO/LRRK2WT
PINK1WT/LRRK2RC
PINK1KO/LRRK2RC
Rab12pSer105
Rab12
merged Rab12
PPM1H
LRRK2pSer935
LRRK2
merged LRRK2
Vinculin
TPS
C.
pRab12
D.
PPM1H
E.
pLRRK2
F.
pRab12
G.
PPM1H
H.
pLRRK2
pS105/Total Rab12 (AU)
PPM1H/Vinculin (AU)
pS935/Total LRRK2 (AU)
Vehicle
MLi2
LRRK2WT
LRRK2RC
PINK1WT
PINK1KO
CEREBELLUM
BRAINSTEM

**Supp4**

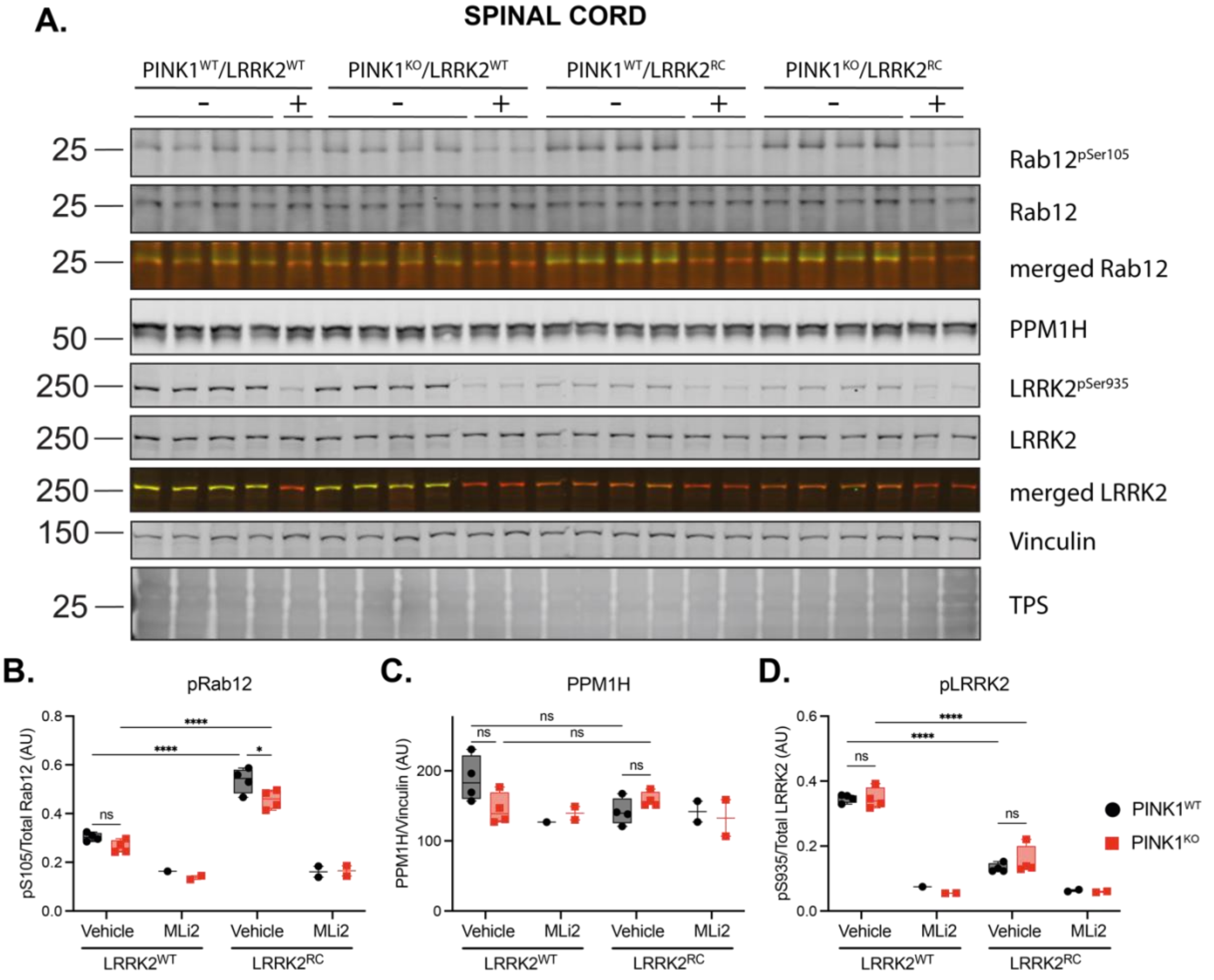

A.
SPINAL CORD
PINK1WT/LRRK2WT
PINK1KO/LRRK2WT
PINK1WT/LRRK2RC
PINK1KO/LRRK2RC
-
+
25
50
250
150
Rab12pSer105
Rab12
merged Rab12
PPM1H
LRRK2pSer935
LRRK2
merged LRRK2
Vinculin
TPS
B.
pRab12
pS105/Total Rab12 (AU)
C.
PPM1H
PPM1H/Vinculin (AU)
D.
pLRRK2
pS935/Total LRRK2 (AU)
ns
****
*
Vehicle
MLi2
LRRK2WT
LRRK2RC
PINK1WT
PINK1KO

### Supp5

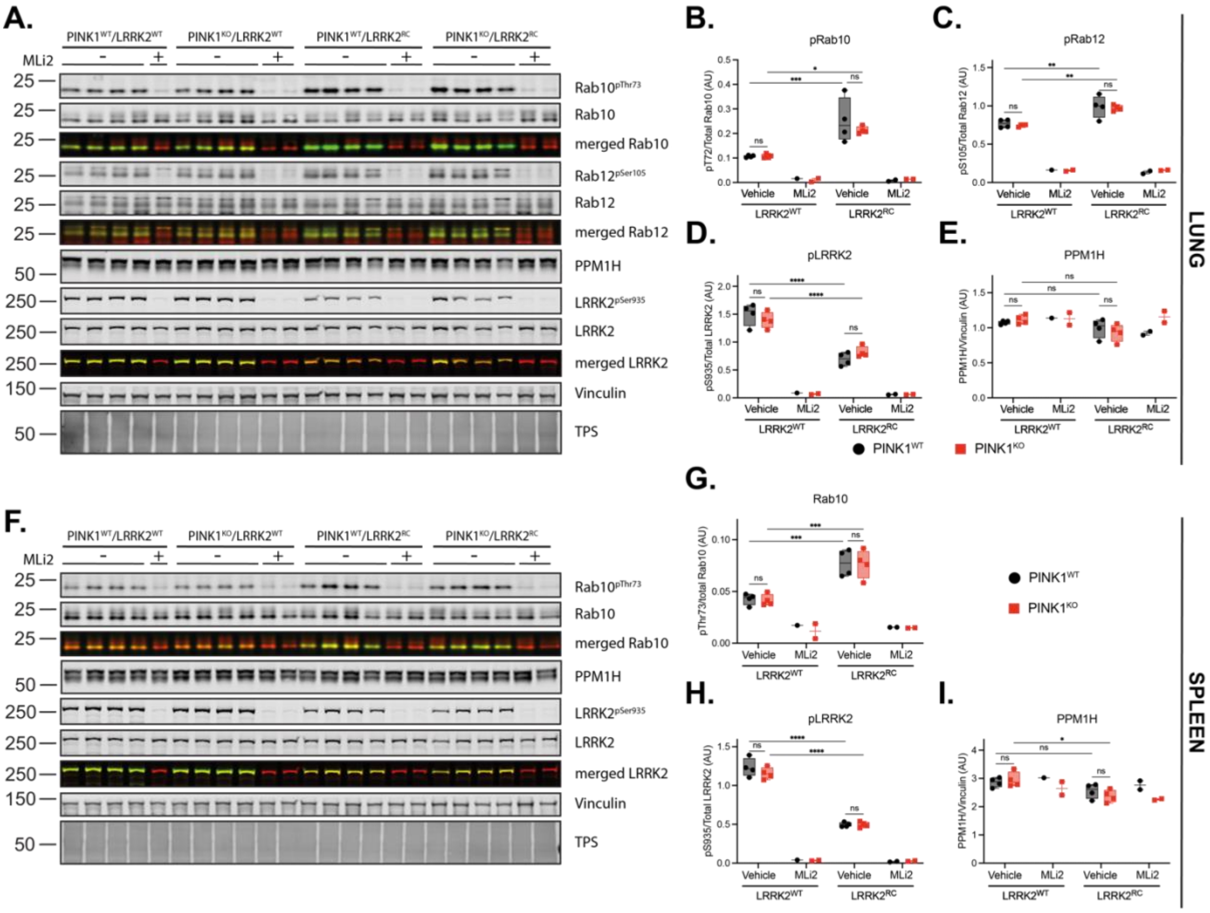

A.
PINK1WT/LRRK2WT
PINK1KO/LRRK2WT
PINK1WT/LRRK2RC
PINK1KO/LRRK2RC
MLi2
Rab10pThr73
Rab10
merged Rab10
Rab12pSer105
Rab12
merged Rab12
PPM1H
LRRK2pSer935
LRRK2
merged LRRK2
Vinculin
TPS
B.
pRab10
pT72/Total Rab10 (AU)
C.
pRab12
pS105/Total Rab12 (AU)
D.
pLRRK2
pS935/Total LRRK2 (AU)
E.
PPM1H
PPM1H/Vinculin (AU)
Vehicle
MLi2
LRRK2WT
LRRK2RC
PINK1WT
PINK1KO
LUNG
F.
G.
Rab10
pThr73/total Rab10 (AU)
H.
pLRRK2
pS935/Total LRRK2 (AU)
I.
PPM1H
PPM1H/Vinculin (AU)
SPLEEN

**Supp6**

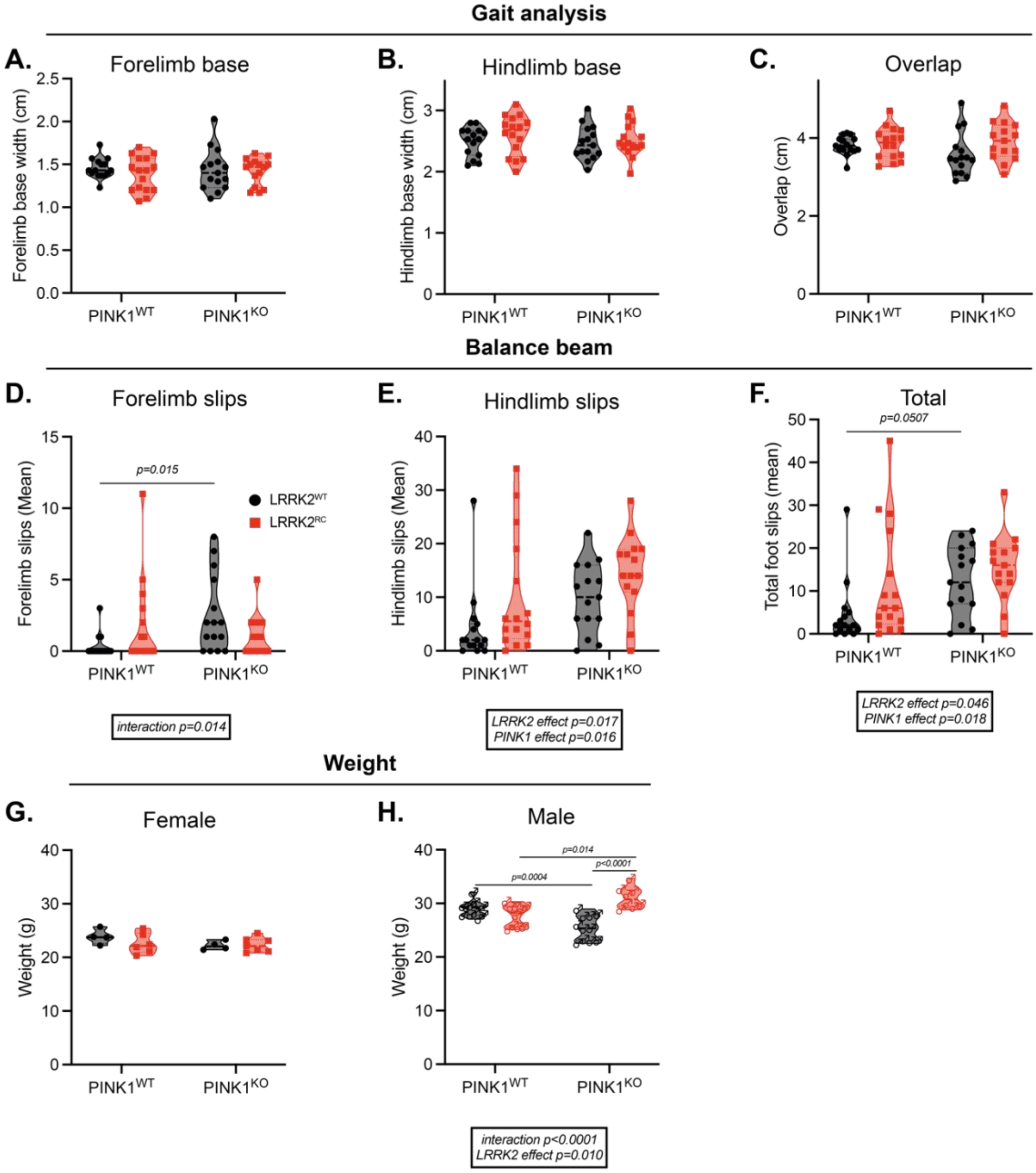
Gait analysis
A.
Forelimb base
Forelimb base width (cm)
2.5
2.0
1.5
1.0
0.5
0.0
PINK1WT
PINK1KO
B.
Hindlimb base
Hindlimb base width (cm)
C.
Overlap
Overlap (cm)
Balance beam
D.
Forelimb slips
Forelimb slips (Mean)
p=0.015
LRRK2WT
LRRK2RC
interaction p=0.014
E.
Hindlimb slips
Hindlimb slips (Mean)
LRRK2 effect p=0.017
PINK1 effect p=0.016
F.
Total
Total foot slips (mean)
p=0.0507
LRRK2 effect p=0.046
PINK1 effect p=0.018
Weight
G.
Female
Weight (g)
H.
Male
Weight (g)
p=0.014
p<0.0001
p=0.0004
interaction p<0.0001
LRRK2 effect p=0.010

Supp7

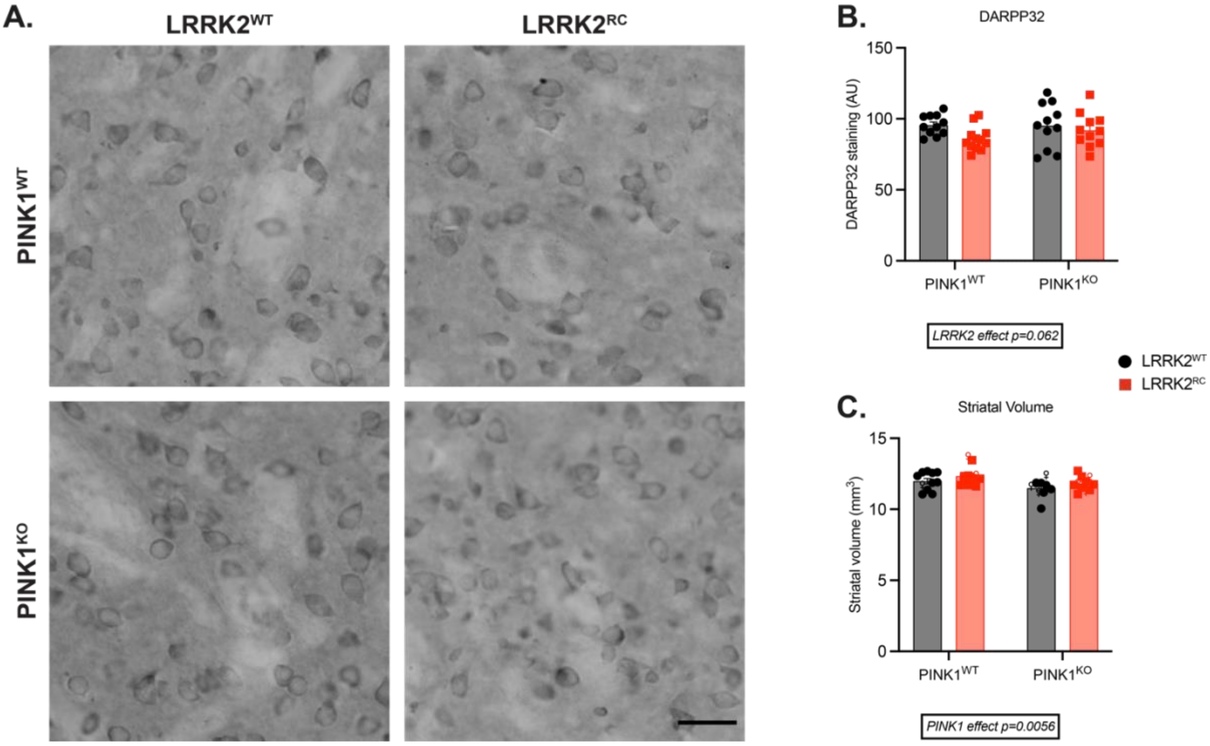

A.
LRRK2$^{WT}$
LRRK2$^{RC}$
PINK1$^{WT}$
PINK1$^{KO}$
B.
DARPP32
DARPP32 staining (AU)
150
100
50
0
PINK1$^{WT}$
PINK1$^{KO}$
LRRK2 effect p=0.062
LRRK2$^{WT}$
LRRK2$^{RC}$
C.
Striatal Volume
Striatal volume (mm$^3$)
15
10
5
0
PINK1$^{WT}$
PINK1$^{KO}$
PINK1 effect p=0.0056

**Supp8**

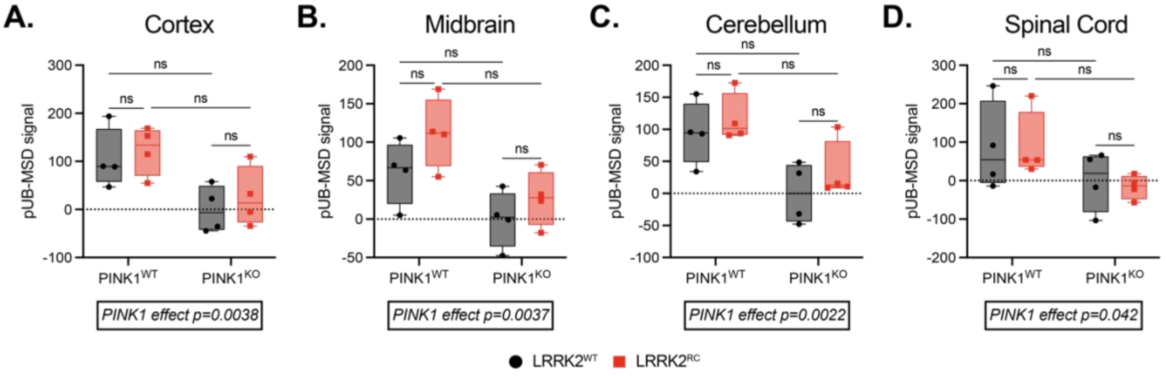
A.
Cortex
B.
Midbrain
C.
Cerebellum
D.
Spinal Cord
pUB-MSD signal
ns
PINK1WT
PINK1KO
PINK1 effect p=0.0038
PINK1 effect p=0.0037
PINK1 effect p=0.0022
PINK1 effect p=0.042
LRRK2WT
LRRK2RC

Supp9

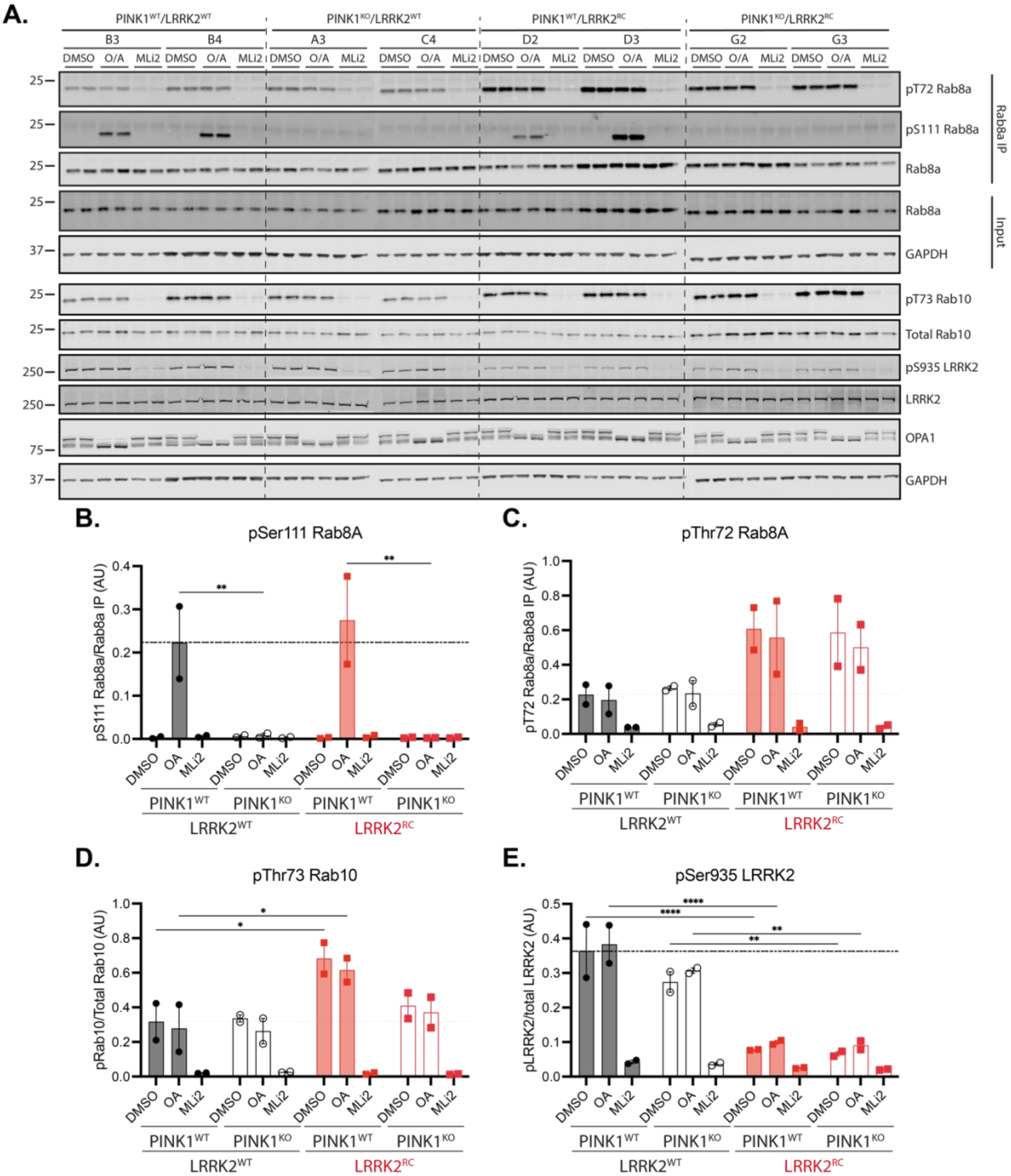
A.
PINK1WT/LRRK2WT
PINK1KO/LRRK2WT
PINK1WT/LRRK2RC
PINK1KO/LRRK2RC
B3
B4
A3
C4
D2
D3
G2
G3
DMSO
O/A
MLi2
pT72 Rab8a
pS111 Rab8a
Rab8a
Rab8a IP
Input
GAPDH
pT73 Rab10
Total Rab10
pS935 LRRK2
LRRK2
OPA1
25
37
250
75
B.
pSer111 Rab8A
pS111 Rab8a/Rab8a IP (AU)
C.
pThr72 Rab8A
pT72 Rab8a/Rab8a IP (AU)
D.
pThr73 Rab10
pRab10/Total Rab10 (AU)
E.
pSer935 LRRK2
pLRRK2/total LRRK2 (AU)
DMSO
OA
MLi2
PINK1WT
PINK1KO
LRRK2WT
LRRK2RC
**
*
****

**Supp10**

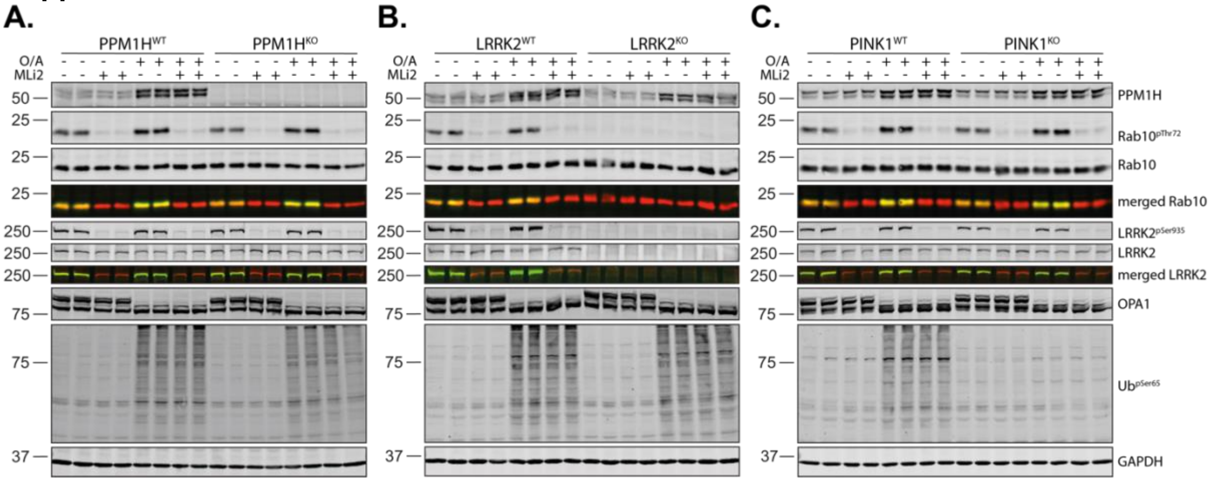
A.
B.
C.
PPM1H^WT
PPM1H^KO
LRRK2^WT
LRRK2^KO
PINK1^WT
PINK1^KO
O/A
MLi2
50
25
250
75
37
PPM1H
Rab10^pThr72
Rab10
merged Rab10
LRRK2^pSer935
LRRK2
merged LRRK2
OPA1
Ub^pSer65
GAPDH

**Supp11**

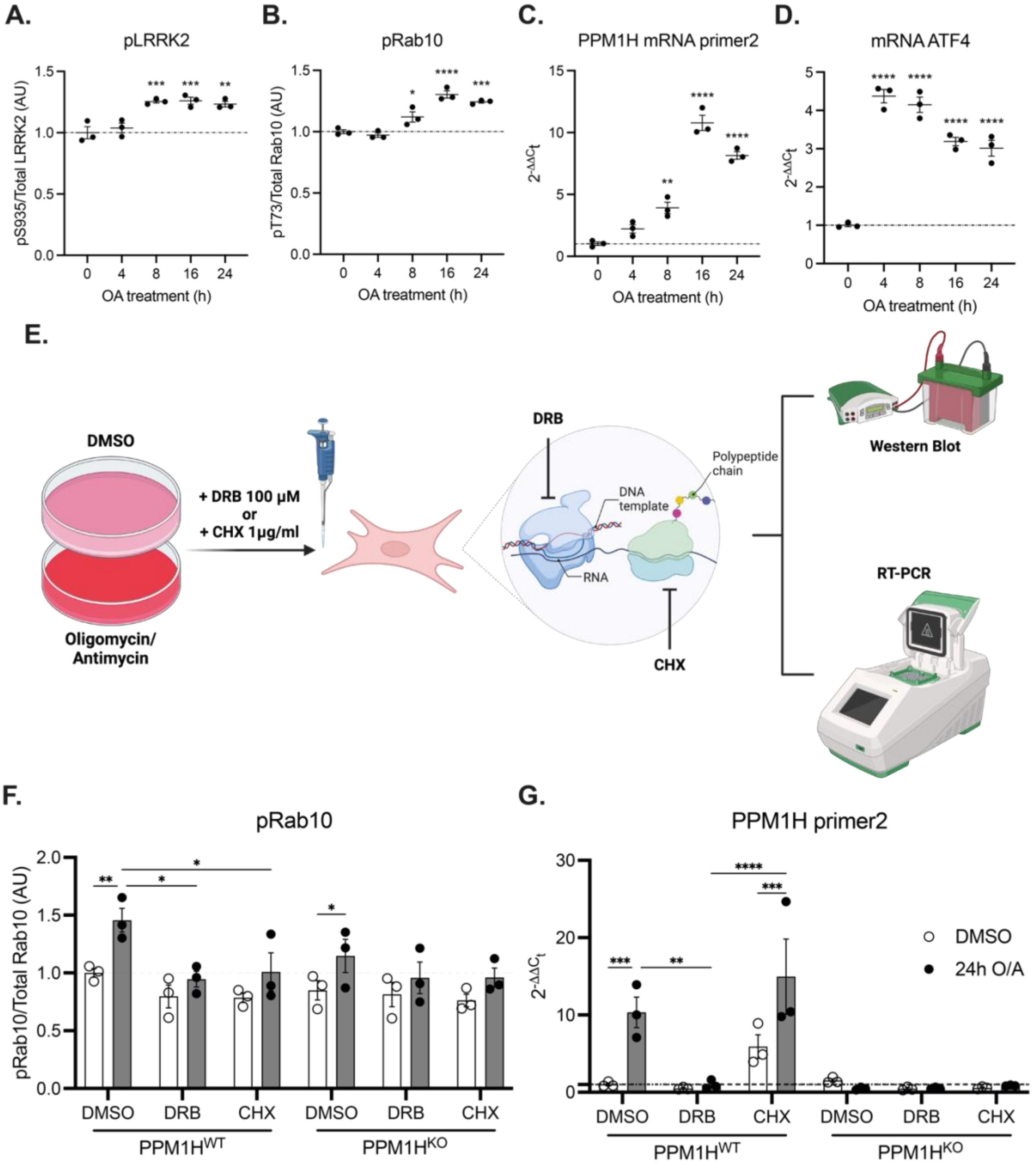

A.
pLRRK2
pS935/Total LRRK2 (AU)
OA treatment (h)
B.
pRab10
pT73/Total Rab10 (AU)
OA treatment (h)
C.
PPM1H mRNA primer2
OA treatment (h)
D.
mRNA ATF4
OA treatment (h)
E.
DMSO
Oligomycin/
Antimycin
+ DRB 100 µM
or
+ CHX 1µg/ml
DRB
DNA template
Polypeptide chain
RNA
CHX
Western Blot
RT-PCR
F.
pRab10
pRab10/Total Rab10 (AU)
DMSO
DRB
CHX
PPM1HWT
PPM1HKO
G.
PPM1H primer2
DMSO
24h O/A
