## Supplementary Key Reagents Table for "Endogenous LRRK2 and PINK1 function in a convergent neuroprotective ciliogenesis pathway in the brain"

| RESOURCE TYPE | RESOURCE NAME | SOURCE | IDENTIFIER | NEW/REUSE | ADDITIONAL INFORMATION |
|---|---|---|---|---|---|
| Antibody | Adenylate cyclase III | EnCor | RPCA-ACIII (AB_2572219 | reuse | 1:10000 |
| Antibody | Arl13b | Neuromab | N295B/66 (AB_2877361) | reuse | 1:1000 |
| Antibody | Biotinylated anti-MOUSE IgG (H+L) | Vector Laboratories | BA-9200 (AB_2336171) | reuse | 1:200 |
| Antibody | Biotinylated anti-rabbit IgG (H+L) | Vector Laboratories | BA-1000 (AB_2313606) | reuse | 1:200 |
| Antibody | Choline Acetyltransferase | Millipore | AB144P-1ML (AB_2079751) | reuse | 1:200 |
| Antibody | CISD1 | CST | 83775S (AB_2800031) | reuse | 1:1000 |
| Antibody | CS | CST | 14309S (AB_2665545) | reuse | 1:1000 |
| Antibody | DARPP32 | Santa Cruz | sc_271111 (AB_10610055) | reuse | 1:100 |
| Antibody | GAPDH | Santa Cruz | Sc-32233 (AB_627679) | reuse | 1:2000 |
| Antibody | GFAP | Encor | CPCA-GFAP (AB_2109953) | reuse | 1:2000 |
| Antibody | H+L Donkey anti-chicken Alexa 488 | Jackson ImmunoResearch | 703-545-155 (AB_2340375) | reuse | 1:2000 |
| Antibody | H+L Donkey anti-goat Alexa 488 | Millipore | A11055 (AB_2534102) | reuse | 1:2000 |
| Antibody | H+L Donkey anti-mouse Alexa 568 | Invitrogen | A10037 (AB_2534013) | | 1:2000 |
| Antibody | H+L Donkey anti-Rabbit Alexa 568 | Millipore | A10042 (AB_2534017) | reuse | 1:2000 |
| Antibody | Iba1 | Wako | 019-19741 (AB_839504) | reuse | 1:1000 |
| Antibody | IRDye 680LT anti-Goat IgG | LI-COR | 926–68 024 (AB_10706168) | reuse | 1:100000 |
| Antibody | IRDye 680LT anti-Mouse IgG | LI-COR | 926–68022 (AB_10715072) | reuse | 1:100000 |
| Antibody | IRDye 680RD anti-Mouse IgG | LI-COR | 926-68070 (AB_10956588) | reuse | 1:100000 |
| Antibody | IRDye 800CW anti-Rabbit IgG | LI-COR | 926-32213 (AB_621848) | reuse | 1:100000 |
| Antibody | IRDye 800CW anti-Rabbit IgG | LI-COR | 926-32211 (AB_621843) | reuse | 1:100000 |

| | | | | | |
|---|---|---|---|---|---|
| Antibody | LAMP1 | CST | 3243S (AB_2134478) | reuse | 1:1000 |
| Antibody | LRRK2 Total C-terminal | Neuromab | N241A/34 (AB_2877351) | reuse | 1 µg/ml |
| Antibody | OPA1 | BD Biosciences | 612606 (AB_399888) | reuse | 1:1000 |
| Antibody | PPM1H | Abcam | ab30353 (AB_2941812) | reuse | 1:1000 |
| Antibody | pSer105 Rab12 | Abcam | ab256487 (AB_2884880) | reuse | 1:1000 |
| Antibody | pSer111 Rab8A | Abcam | ab267492 (AB_3099419) | reuse | 1:1000 |
| Antibody | pSer2 RNA polymerase | Abcam | ab5095 (AB_304749) | reuse | 1:5000 |
| Antibody | pSer65 Ubiquitin | CST | 62802 (AB_2799632) | reuse | 1:1000 |
| Antibody | pSer935 LRRK2 | MRC PPU Reagents and services, UoD | UDD2 10(12) (AB_2921228) | reuse | 1 µg/ml |
| Antibody | pThr72 Rab8a | Abcam | ab230260 (AB_2814988) | reuse | 1:1000 |
| Antibody | pThr73 Rab10 | Abcam | ab230261 (AB_2811274) | reuse | 1:1000 |
| Antibody | Rab10 Total | Nanotools | 0680-100/Rab10-605B11 (AB_2921226) | reuse | 1 µg/ml |
| Antibody | Rab12 Total | MRC PPU Reagents and services, UoD | SA227 (AB_2921227) | reuse | 1 µg/ml |
| Antibody | Rab8A total | Abcam | ab237702 (AB_3099418) | reuse | 1:1000 |
| Antibody | TOM20 | Abcam | ab186735 (AB_2889972) | reuse | 1:1000 |
| Antibody | Tubulin | Proteintech | 66031-1-IG (AB_11042766) | reuse | 1:10000 |
| Antibody | Ubiquitin | Biolegend | 646302 (AB_1659270) | reuse | 1:1000 |
| Antibody | Vinculin | Abcam | ab129002 (AB_11144129) | reuse | 1:10000 |
| Chemical | Antimycin | Sigma-Aldrich | A8674 | reuse | |

| | | | | | |
|---|---|---|---|---|---|
| Chemical | CHX, cycloheximide | Sigma-Aldrich | C7698 | reuse | |
| Chemical | DMSO | Sigma-Aldrich | D2650 | reuse | |
| Chemical | DRB, 5,6-Dichlorobenzimidazole 1-β-D-ribofuranoside | Sigma-Aldrich | D1916 | reuse | |
| Chemical | Fast green FCF | Sigma-Aldrich | F7252 | reuse | for total protein staining |
| Chemical | Mli-2 | MRC PPU Reagents and services, UoD | | reuse | |
| Chemical | Oligomycin | Sigma-Aldrich | 75351 | reuse | |
| Chemical | PowerUp™ SYBR™ Green Master Mix for qPCR | Thermo Fisher | A25742 | | |
| Chemical: primer | mouse ACTB FW | Sigma-Aldrich | TGACGTTGACATCCGTAAAG | reuse | PureSimplePrimer |
| Chemical: primer | mouse ACTB REV | Sigma-Aldrich | GAGGAGCAATGATCTTGATCT | reuse | PureSimplePrimer |
| Chemical: primer | mouse GAPDH FW | Sigma-Aldrich | TGACCTCAACTACATGGTCTACA | reuse | PureSimplePrimer |
| Chemical: primer | mouse GAPDH REV | Sigma-Aldrich | CTTCCCATTCTCGGCCTTG | reuse | PureSimplePrimer |
| Chemical: primer | mouse PPM1H FW1 | Sigma-Aldrich | ATATGGAGAAGGCAAGAAGG | reuse | PureSimplePrimer |
| Chemical: primer | mouse PPM1H FW2 | Sigma-Aldrich | AAAACCATTCCTGTCTTCAG | reuse | PureSimplePrimer |
| Chemical: primer | mouse PPM1H REV1 | Sigma-Aldrich | TCATATCTGGAGAGATCGTAG | reuse | PureSimplePrimer |

| | | | | | |
|---|---|---|---|---|---|
| Chemical: primer | mouse PPM1H REV2 | Sigma-Aldrich | TCTGGATCACAGTTAGGAAG | reuse | PureSimplePrimer |
| Chemical: primer | mouse Rpl13a FW | Sigma-Aldrich | AGCCTACCAGAAAGTTTGCTTAC | reuse | PureSimplePrimer |
| Chemical: primer | mouse Rpl13a REV | Sigma-Aldrich | GCTTCTTCTTCCGATAGTGCATC | reuse | PureSimplePrimer |
| Chemical: primer | mouse TBP FW | Sigma-Aldrich | CCTTGTACCCTTCACCAATGAC | reuse | PureSimplePrimer |
| Chemical: primer | mouse TBP REV | Sigma-Aldrich | ACAGCCAAGATTCACGGTAGA | reuse | PureSimplePrimer |
| Dataset | Figure 1, figure 3 and figure 4 | Zenodo | 10.5281/zenodo.11244913 | New | All data presented in Figure 1, figure 3 and figure 4 |
| Dataset | Figure 2 | Zenodo | 10.5281/zenodo.11518850 | | All raw data presented in Figure 2 |
| Dataset | Figure 5 | Zenodo | 10.5281/zenodo.11519152 | | All raw data presented in Figure 5 |
| Dataset | Supplementary Figures | Zenodo | 10.5281/zenodo.11546690 | | All raw data presented in Supplementary |
| Experimental model:organism | LRRK2 WT and LRRK2 KO | Jax | IMSR_JAX:012453 | reuse | Used for mouse embryonic fibroblasts |
| Experimental model:organism | LRRK2 WT and R1441C mice | Jax | IMSR_JAX:009346 | reuse | Used for mouse embryonic fibroblasts |
| Experimental model:organism | PINK1 WT and PINK1 KO mice | Jax | | reuse | Used for mouse embryonic fibroblasts |

| | | | | | |
|---|---|---|---|---|---|
| Experimental model:organism | PINK1 WT/LRRK2 WT, PINK1 WT/LRRK2 RC, PINK1 KO/LRRK2 WT, PINK1 KO/LRRK2 RC | Double mutants generated by crossing the two lines | | new | In vivo experiments and generation of mouse embryonic fibroblasts |
| Experimental model:organism | PPM1H WT and PPM1H KO | Taconic | TF3142 | reuse | Used for mouse embryonic fibroblasts |
| Kit | PureLink™ RNA Mini Kit | Thermo Fisher | 12183018A | reuse | |
| Kit | QuantiTect Reverse Transcription Kit | Qiagen | 205311 | reuse | |
| Kit | RNAscope multiplex fluorescent reagent kit | Advanced Cell Diagnostics | 323100 | reuse | |
| Kit | RNAscope Probe-Mm-*Gdnf*-C1 | Advanced Cell Diagnostics | 421951 | reuse | 1:20 dilution |
| Protocol | Cell and tissue immunoblotting | protocols.io | dx.doi.org/10.17504/protocols.io.ewov14znkvr2/v2 | reuse | |
| Protocol | MEF generation and mainteinance | protocols.io | dx.doi.org/10.17504/protocols.io.eq2ly713qlx9/v1 | reuse | |
| Protocol | Mice behavioural test | protocols.io | | reuse | |
| Protocol | Mitochondrial fractionation | protocols.io | dx.doi.org/10.17504/protocols.io.bxmypk7w | reuse | |
| Protocol | PINK1 siRNA | protocols.io | dx.doi.org/10.17504/protocols.io.kxygx343zg8j/v1 | new | |
| Protocol | Primary cilia analysis | protocols.io | dx.doi.org/10.17504/protocols.io.bnwimfce | reuse | |
| Protocol | pSer65 Ub ELISA | method article | doi: 10.1080/15548627.2020.1834712. | reuse | |

| | | | | |
|---|---|---|---|---|
| Protocol | RT-PCR | protocols.io | dx.doi.org/10.17504/protocols.io.81wgbz7r3gpk/v1 | new |
| Protocol | Subcutaneus injection | protocols.io | dx.doi.org/10.17504/protocols.io.bezdjf26 | reuse |
| Protocol | Fluorescence in situ hybridization (FISH) | bioprotoccol | bio-protocol.org/prep1423 | reuse |
| Software | FiJI | ImageJ | SCR_002285 | reuse |
| Software | Graphpad | Prism | SCR_002798 | reuse |
| Software | ImageStudio | Biorad | SCR_013715 | reuse |
